# Ecological assembly processes of the bacterial and fungal microbiota of wild and domesticated wheat species

**DOI:** 10.1101/2020.01.07.896910

**Authors:** M. Amine Hassani, Ezgi Özkurt, Sören Franzenburg, Eva H. Stukenbrock

## Abstract

Plant domestication has led to substantial changes in the host physiology. How this anthropogenic intervention has contributed in altering the wheat microbiota is not well understood. Here, we investigated the role of ecological selection, drift and dispersal in shaping the bacterial and fungal communities associated with domesticated wheat *Triticum aestivum* and two wild relatives, *Triticum boeoticum* and *Triticum urartu*. Our study shows that the bacterial and fungal microbiota of wild and domesticated wheat species follow distinct community assembly patterns. Further, we revealed a more prominent role of neutral processes in the assembly of the microbiota of domesticated wheat and propose that domestication has relaxed selective processes in the assembly of the wheat microbiota.

---

Early crop cultivation and domestication were key in the rise of agrarian societies and civilizations (Diamond 2002). Wheat represents one of the first cereal crop species, domesticated 10,000 years ago in the Fertile Crescent, it underwent through a reticulated evolution (Pont et al. 2019). The (un)conscious anthropogenic selection led to substantial alterations of the plant physiology which are hallmarked by seed size enlargement (Purugganan et al. 2009), loss of shattering (Fuller et al. 2007) and increased yield (Preece et al. 2017). Archaeobotanical studies evidenced that the spread of cereal crops from their center of origin through human migration and trade further contributed to their diversification (Purugganan et al. 2009). Concomitantly, these multi-step and sequential processes caused genetic bottlenecks and reduced genetic diversity in the genome of domesticated crops (Doebley et al. 2006).

Investigating domesticated plant species has been key to comprehend evolutionary processes that contributed to plant traits emergence (Purugganan et al. 2009) and their molecular genetic basis (Doebley et al. 2006). However, our understanding of the impact of domestication on the plant associated microbiota remains incipient. By comparing wild and modern crop genotypes, few studies have addressed the effect of domestication in altering the composition and structure of the plant-associated bacterial and fungal microbiota, (recently reviewed in Hassani et al. 2019; Escudero-Martinez et al. 2019; Cordovez et al. 2019). Depending on the crop species, plant domestication has resulted in microbial diversity reduction (Zachow et al. 2014), reduced symbiotic interactions (Kiers et al. 2007) or depletion of a microbial phylogroup (Pérez-Jaramillo et al. 2018). While genetic diversity of crop species is reduced at a population level, we still lack insight why individual crop plants assemble less diverse microbial communities compared to their wild relatives. Furthermore, it remains unclear to what extent plant domestication has altered ecological processes (i.e. selection, drift and dispersal) of the microbiota assembly. Here, we propose that wild and domestication wheat species show distinct assembly patterns of bacterial and fungal microbiota and that selection processes are less eminent in the assembly of the domesticated wheat *Triticum aestivum*. To characterize the role of ecological drift, dispersal and selection in governing the assembly of the wheat microbiota, we studied the bacterial and fungal microbiota of the domesticated wheat species *T. aestivum* and two of its wild relatives (i.e. *Triticum boeoticum* and *Triticum urartu*). In our analyses and interpretations, we followed the conceptual framework of microbial community assembly that pose ecological drift, selection and dispersal as processes acting in concert, and not in disjoint, to drive microbial patterns (Nemergut et al. 2013; Paredes et al. 2016; Vellend et al. 2014).

To compare the microbiota between wheat species, *T. boeoticum*, *T. urartu* and *T. aestivum* were grown in an agricultural soil under controlled growth conditions. After 10 days, bacterial and fungal communities associated with leaves, roots and rhizosphere of each individual plant, as well as unplanted soil, were profiled by the sequencing of the microbial maker genes 16S and ITS (see Methods). The analysis of the community composition indicated that leaf and rhizosphere habitats of *T. urartu* harbor more diverse bacteria than *T. boeoticum* and *T. aestivum* (Supplementary Fig. 1a). Furthermore, the communities associated with *T. urartu* showed a trend towards an increased evenness (higher Shannon index - Supplementary Fig. 1b) and phylogenetic breadth (Supplementary Fig. 1c). In contrary, only the fungal root microbiota of *T. urartu* showed an increase in species richness and evenness (Supplementary Fig. 1d-e). Several bacteria belonging to Acidobacteria, Firmicutes and Proteobacteria mediated the higher alpha-diversity observed in the rhizosphere of *T. urartu* (Supplementary Fig. 2), whereas the increased fungal diversity in the roots of *T. urartu* was explained by Ascomycota, Basidiomycota, Mortierellomycota and other fungal phyla (Supplementary Fig. 3). The finding that *T. boeoticum* and *T. aestivum* harbor comparably diverse microbiota indicates that wheat domestication did not systematically impair microbial diversity and reveals that microbial richness can differ even among wild species.

To further address how plant habitat (i.e. leaf, root or rhizosphere) and host genetics shape the structure of the microbiota, we projected Bray-Curtis (BC) distances between all samples using an ordination method (see Methods). Plant habitat and second wheat genetic identity had significant effect on the assembly of the bacterial and fungal communities (Supplementary Fig. 4 and 5, respectively). Remarkably, the structure of the bacterial communities associated with the rhizosphere of *T. boeoticum* was more distinct from those of *T. urartu* than *T. aestivum* (Supplementary Fig. 6a and b). The corresponding rhizosphere fungal microbiota of these wheat species did not follow similar patterns (Supplementary Fig. 6c). These results suggest that root exudations of *T. urartu* to a higher extent favor the enrichment of bacteria. Whether this phenotype is mediated by an increased exudation of “simple” sugars by *T. boeoticum* as in *T. aestivum* (Shaposhnikov et al. 2016) remains hypothetical and will require future testing.

To reveal bacterial or fungal taxa that contribute to the observed differences in the community structure, we tested the enrichment of bacterial or fungal OTUs in one wheat species compared to another (see Methods). We show that the observed difference in the community structure of bacteria across the three wheat species is explained by the enrichment of several OTUs (Supplementary Fig. 7-9), these OTUs belong mainly to Proteobacteria, Actinobacteria, Firmicutes and Bacteroidetes (Supplementary Fig. 10). Differences in the structure of fungal communities were less pronounced (Supplementary Fig. 11-13). In contrast to bacteria (Supplementary Fig. 10), no single fungal taxa, among the significantly enriched OTUs, was shared between root and rhizosphere habitats (Supplementary Fig. 14). These data indicate that fungi, unlike bacteria, do not follow the multi-step microbiota establishment (Edwards et al. 2015; Bulgarelli et al. 2013) and point to a more pronounced role of dispersal in the assembly of the bacterial microbiota compared to the fungi. Taken together, these data indicate that plant habitat and host genetics are selective factors that govern the assembly of the bacterial and fungal microbiota of wild and domesticated wheat species. The interaction of these two factors contributes to the assembly of distinct microbial communities.

Wild and domesticated wheat species assemble distinct bacterial and fungal communities. These communities show different degrees of variance in their structure (Özkurt et al. 2019). To compare community structure homogeneity across the microbiota of wild and domesticated wheat species, we computed BC (for bacteria and fungi) and weighted UniFrac (only for bacteria) distances to the centroid (see Methods). Bacterial and fungal communities associated to the leaves of *T. urartu* showed less heterogeneity than communities of *T. boeoticum* and *T. aestivum* (Fig. 1a-b). Additionally, leaf-associated bacteria of both *T. boeoticum* and *T. urartu* showed increased phylogenetic clustering compared to the leaf-associated bacteria of *T. aestivum* (Fig. 1b). These data indicate that the wild wheat *T. urartu* tends to assemble more homogeneous bacterial and leaf-associated fungal communities. The current analysis also reveals a substantial relative contribution of stochastic processes in the assembly of the microbiota.

**Figure 1.**
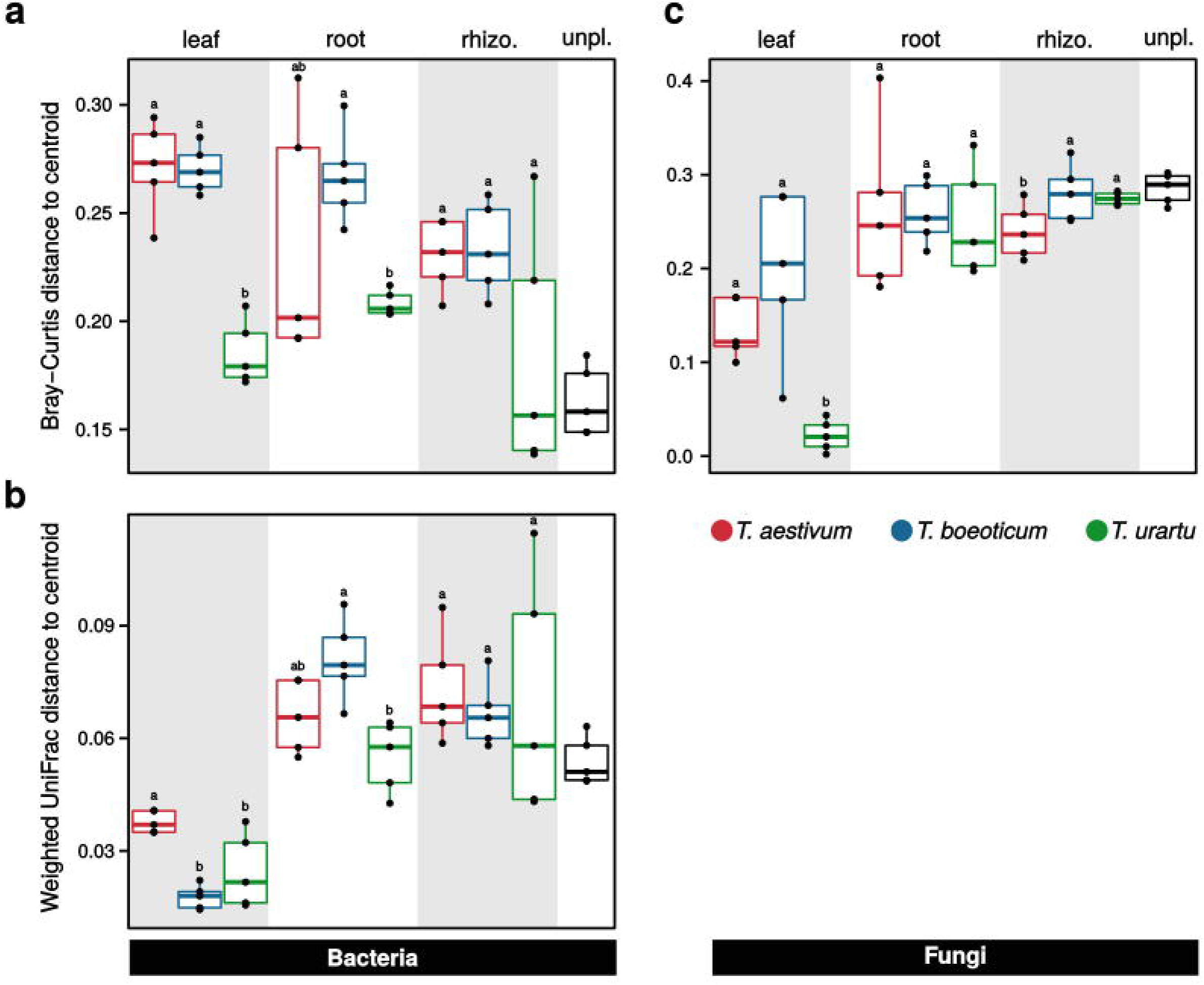
Heterogeneity in structure of bacterial and fungal wheat microbiota. Panels **a** and **b** show community homogeneity analyses depicted by Bray-Curtis and weighted UniFrac distances to centroid for the bacterial microbiota, respectively. **a**, the decrease in BC distance to centroid indicates that the leaf and root microbiota of the wild wheat *T. urartu* assemble more homogeneous bacterial communities than *T. boeoticum* and *T. aestivum*. **b**, both leaf communities associated to *T. boeoticum* and *T. urartu* are more phylogenetically clustered than bacterial communities associated to *T. aestivum*. **c**, across root and rhizosphere habitat, wheat genotype do not explain fungal community heterogeneity. Leaf fungal microbiota of *T. urartu* shows an increased homogeneity than the fungal microbiota associated to *T. boeoticum* and *T. aestivum*. Count data were normalized using cumulative sum scaling factor and OTUs with an occurrence <3 were trimmed out prior to computing distances. Differences in group means was tested using Tukey’s Honest test (0.95 confidence level).

To assess the relative contribution of stochasticity in the assembly of the microbiota on wild and domesticated wheat, we fitted bacterial and fungal data to the Sloan neutral (Sloan et al. 2006) and beta-null (Trucker et al. 2016) models, respectively (see Methods). By defining soil bacteria as seed-source (Venkataraman et al. 2015), we showed that the assembly of the bacterial wheat microbiota do not, or very poorly, follow neutral processes (Supplementary-table 1). This finding additionally corroborates the role of selection in structuring microbial communities. Next, we constrained our analysis to within-group bacterial communities (i.e. across replicates of a same wheat species and corresponding plant habitat, soil bacteria were not defined as seed-source) and quantified ecological drift (*R*^2^ - value) and dispersal (computed as immigration coefficient to the model, *m* – value). Except for the phyllosphere of *T. aestivum*, the fit values to the neutral model were overall higher in the unplanted soil and rhizosphere compared to the root and leaf bacterial communities (Fig. 2a – x-axis, Supplementary-table 2 – *R*^2^ value). Similarly, unplanted soil and rhizosphere habitats exhibited higher immigration coefficients (Fig. 2a – y-axis, Supplementary-table 2 – *m* value). By comparing the fit and immigration values to the model across plant habitats, we conclude that root- and leaf-associated bacteria are more prone to selection than rhizosphere communities which suggests that additional plant-related, and possible un-related, cues contribute to these processes. The relative contribution of neutral processes in the assembly of the microbiota was not only depending on plant habitat, but was also reflected by the wheat genetic identity. Remarkably, neutral processes are better explaining the assembly of the root and leaf microbiota of the domesticated wheat *T. aestivum* than of the two wild relatives (Fig. 2a – x-axis, Supplementary-table 2 – R^2^ value). The explanation of these findings resides on the fact that fewer bacterial OTUs deviated from neutral expectations in *T. aestivum* (Supplementary Fig. 15 - unfilled circles), whereas more OTUs were reported above or below neutral prediction (i.e. over- or under-represented, respectively) in both *T. boeoticum* (Supplementary Fig. 15) and *T. urartu* (Supplementary Fig. 15). Taken together, this analysis indicates that neutral processes have a more prominent role in the assembly of the root and leaf bacterial microbiota of the domesticated wheat *T. aestivum* compared to its wild relatives.

**Figure 2.**
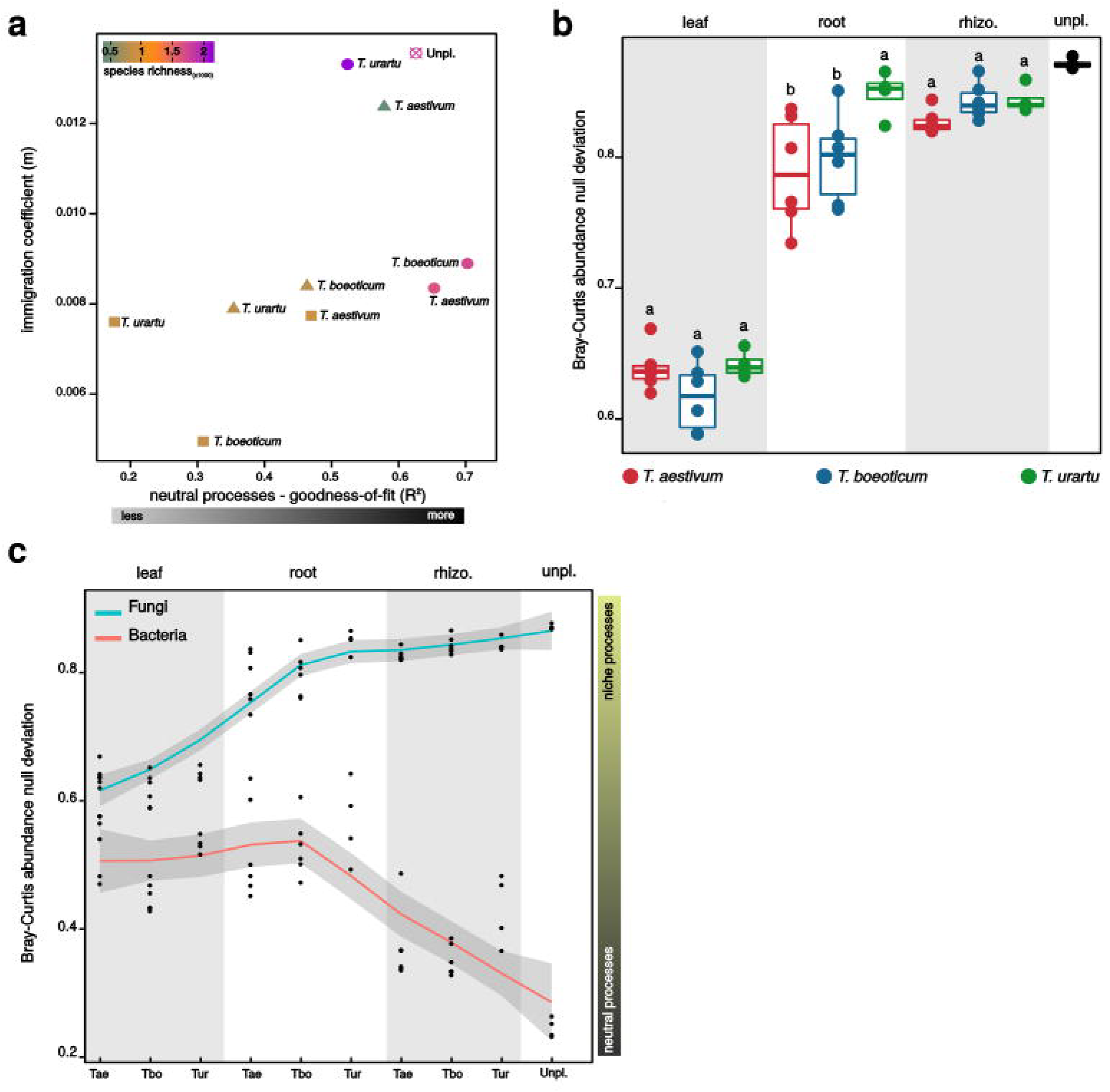
Ecological assembly processes of the bacterial and fungal microbiota of wheat. **a**, the scatter-plot depicts the goodness of fit (*R*^*2*^) and immigration coefficient (*m*) to the neutral model of bacterial communities. In the graph, square, triangle and circle indicate root, leaf and rhizosphere bacterial communities, respectively. Unplanted soil is indicated by a crossed unfilled circle. Color gradient in the shapes indicates species richness. The root and leaf microbiota of wild wheat show decreased fit to the neutral processes. The root and leaf bacterial microbiota of wild wheat show a lower fit to the neutral model than *T. aestivum*. **b**, box-plot depicts Bray-Curtis deviation from the null model of the fungal microbiota of wheat across leaf, root and rhizosphere habitats. The root fungal microbiota of *T. boeoticum* and *T. aestivum* show further deviation from niche processes than the fungal microbiota of *T. urartu*. Significance within-group medians were tested using Kruskal-Wallis significance and Conover’s multiple comparisons tests. **c**, The plot depicts Bray-Curtis deviation from the null model of the bacterial and fungal microbiota of wheat across leaf, root and rhizosphere habitats. Color in the line indicates the microbial kingdom. Lines show local regression fitting (Loess, x∼y, 0.95 confidence interval) in the BC-null deviations. Singletons were trimmed out from the count data prior to the model computation.

To quantify the relative contribution of niche and neutral processes in the assembly of the fungal microbiota of wheat, we opted to compute Bray-Curtis beta-diversity deviations from null expectation (Tucker et al. 2016; Lee et al. 2017). Irrespective of the wheat species, fungal communities associated to root and rhizosphere deviated further from null expectations than the leaf fungal microbiota that fell closer to niche processes (Fig 2b). Across wheat species, neither leaf nor rhizosphere fungal microbiota showed a signification difference in the beta-null deviation. However, the assembly of fungal communities on the roots of *T. urartu* was more prone to niche processes compared to *T. boeoticum* and *T. aestivum*. The analysis of BC deviation from null model indicates that the transition from the rhizosphere through the root to the leaf habitat is concomitant with a decrease in niche processes. Remarkably, similar analysis conducted on the bacterial communities showed the opposite, i.e. an increase in niche processes (Fig 2c). We conclude from these results that the assembly of bacterial and fungal communities follow distinct community patterns and further suggest that wheat domestication and breeding have a lesser extent altered the assembly of the wheat fungal microbiota. The divergent patterns are most likely mediated by the constrained dispersal of fungi (Supplementary fig. 16) and the scarcity of dominant fungal OTUs across individual plants (Supplementary fig. 17).

In this study, we have investigated bacterial and fungal communities that assemble on wild and domesticated wheat species. The analysis of the community structure indicated that plant habitat and host genetics both contribute in shaping bacterial and fungal communities. Microbial richness between wild and domesticated wheat species was mitigated across habitats and microbial kingdoms, with a tendency of the wild wheat *T. urartu* to harbor more diverse microbiota. By deciphering ecological processes in the assembly of the microbiota, we showed that neutral processes have a more prominent role in the assembly of the bacterial microbiota of *T. aestivum*. Based on these findings we suggest that domestication might have relaxed selective processes that govern the assembly of the microbiota in the root and leaf habitats of *T. aestivum*. Whether this phenotype is related to a loss of genetic cues in the modern wheat will require future testing.

## Methods

### Soil sample and wheat accessions

The soil was collected during summer 2017 from the experimental farm Hohenschulen (field Scheunenkopp, 54°18’53.7"N – 9°58’44.1"E) of the Christian-Alberchts University of Kiel, Germany. After clearing off the surface, soil was taken from the top layer (∼20 cm). The soil was homogenized, passed through 5 mm sieve, spiked with 5% (v/v) of peat (HAWITA GmbH, Germany) and stored at room temperature for 3 additional months. The domesticated wheat, *Triticum aestivum*, used in this study, is a landrace that originates from the province of Kislak Antakya in Turkey (35° 58’ 21.8’’N – 36° 07’ 41.8"E). Wild wheat varieties (i.e. *T. boeoticum* - TRI 18344 and *T. urartu* - TRI 18414) originate from Turkey and were kindly provided by the IPK Gatersleben, Germany.

### Growth conditions

Prior to sow the seeds, these were surface sterilized through three successive washing steps as indicated in the following. Seeds were washed (1) 3x with ddH_2_O (Veolia water technologies), (2) 1x with sterile 1% phosphate-buffered saline (v/v, PBS) supplemented with 0.02 % (v/v) Tween 20, (3) 3x ddH_2_O, then (4) 1x with 80% (v/v) ethanol and rinsed (5) 3x with ddH_2_O. All steps were carried out using sterile 50 ml sterile tubes (Sarstedt AG & Co, Germany). Each individual seed was sowed in square PVC pots containing 415 g of Kiel agricultural soil. All pots containing the same wheat genotype were placed in the same tray and incubated for additional 10 days in a phytochamber with 16/8 hours light [∼200 µmol/m^2^ 355/s]/dark cycle, 20° C and 90% relative humidity as growth conditions. Plants were watered with sterile tap water every other day by tray flooding.

### Leaf, root and rhizosphere sampling

Leaf, root and rhizosphere samples were harvested for each individual plant separately and kept so for all processes from DNA extraction to PCR-amplicons sequencing. The 1^st^ leaf (7 cm) for each plant was cut and subjected to successive washing using 20 ml of (1) ddH_2_O, (2) 1% (v/v) PBS suppl. with 0.02% (v/v) Tween 20 and (3) 1% BPS. Leaves were briefly dried using sterile blotting paper (0.35 mm, Hahnemühle, Germany). Remaining plant was gently dug out from soil-containing pot and shaken by hands to remove loosely attached soil. Below-ground tissue was separated from the above part, placed in 50 ml sterile tube (Sarstedt AG & Co, Germany) and 20 ml of 1% BPS was added to it. To recover the rhizosphere, the tube was mixed by vortex during 30 sec at max speed. Recovered rhizosphere soil was resuspended in 3 ml sterile ddH_2_O. Corresponding roots were additionally washed following same procedures as indicated above. Each of leaf, root and rhizosphere samples were transferred to a separate Lysing Matrix E tubes (MP Biomedicals, Santa Ana), snap frozen in liquid nitrogen and then stored at −80° C. To maximize recovery of DNA yield, each sample was separated into three subsamples prior to their transfer to lysing tubes.

### DNA extraction

Recovered samples were homogenized twice by Precellys 24 TissueLyser [6300 rpm/15 sec/10 sec pause] (Bertin Technologies, Germany). The DNA extractions were conducted according to the manufacturer’s protocol (FastDNA SPIN kit for Soil; MP Biomedicals, USA), except for two additional steps that were included after homogenization. Samples were pre-treated with (1) 36 µl of Lysozyme [1% w/v in 1% PBS – 37°C/30min with intermittent shaking] (Merck KGaA, Germany) and then with (2) 15 µl Proteinase K [>600 mA/ml] and 5 µl Rnase A [100 mg/ml] (Qiagen, Germany) for 5min at room temperature. DNA was eluted in nuclease-free water (Merck KGaA, Germany) and stored at −20°C for downstream processing.

### Bacterial and fungal community profiling

To profile bacterial and fungal microbiota, 16S rRNA gene and ITS DNA library for Illumina sequencing were prepared through two-step PCR-amplification protocol. Briefly, DNA concentration for each sample was adjusted to 50 ng/µl (NanoDrop 2000c, Thermo Fisher Scientific). Each sample was amplified two independent times and kept so during whole procedure indicated in the following. Bacterial DNA was amplified using the forward primer 799F [5’-AACMGGATTAGATACCCKG-3’] and reverse primer 1192R [5’-ACGTCATCCCCACCTTCC-3’]. To block plant mitochondrial DNA, ten times the volume 799F or 1193R of the blocking primer [5’-GGCAAGTGTTCTTCGGA/3SpC3/-3’] was added to each reaction. Fungal DNA was amplified using the primers set ITS1F [5’-CTTGGTCATTTAGAGGAAGTAA-3’] and ITS1R [5’-GCTGCGTTCTTCATCGATGC-3’]. Both 16S rDNA and ITS regions were amplified by PCR in triplicates [95°C/2min - 95°C/30sec, 55°C/30sec, 72°C/45sec for 25 cycle - 72°C/10min]. Technical replicates were pooled and leftover primers were enzymatically digested for 30min at 37°C by adding 3 µl of Exonuclease I [20 U], 3 µl Antarctic phosphatase [5 U] and 7.32 µl Antarctic phosphatase buffer (both New England BioLabs, Germany). The reaction was heat-inactivated at 85°C during 15 min. A unique bar-code sequence (reverse Illumina compatible primers B1 to B120 and F1 to F120 for bacteria and fungi, respectively) was added to the amplicons obtained from the 1^st^ PCR-reactions over 10 PCR-cycles [95°C/2min - 95°C/30sec, 55°C/30sec, 72°C/45sec - 72°C/10min]. All PCR reactions were performed using Q5^©^ high-fidelity DNA polymerase [2,000 units/ml] (New England Biolabs, Germany) in triplicates. For 16S rDNA amplicons, PCR replicates from 2^nd^ reaction were pooled and run through 1.5% (w/v) agarose gel [90V for ca. 2h-30min] and a band with size ca. 0.3 kb was extracted using QIAquick Gel Extraction Kit (Qiagen, Germany) according to the manufacturer’s protocol. For ITS amplicons, PCR replicates were pooled and remaining primers were enzymatically digested as indicated above and purified using QIAquick PCR Purification Kit (Qiagen, Germany). PCR-products were quantified (NanoDrop 2000c, Thermo Fisher Scientific) and concentration adjusted to 20ng/µl prior to be pooled. Bacterial and fungal PCR-amplicon libraries were cleaned twice with Agencourt AMPure XP Kit (Beckman Coulter, Germany) and submitted for DNA sequencing using the MiSeq Reagent kit v3 with the 2 × 300 bp paired-end sequencing protocol (Illumina Inc., USA).

### Sequence processing and community analysis

Forward and reverse reads were joined, demultiplexed using Qiime2 pipeline (2018.11.0) (Bolyen et al. 2019). PhiX and chimeric sequences were filtered out using Qiime2-DADA2 (Callahan et al. 2016). Scripts used for sequence processing are available under [https://github.com/hmamine/MWDW/tree/master/reads_preprocessing] and raw MiSeq raw read are accessible using the following link [https://www.ncbi.nlm.nih.gov/sra/PRJNA590366]. For Shannon and Observed OTUs analyses, count reads were rarefied to an even sequencing depth based on the smallest sample size for bacterial communities and not rarefied for fungal communities using the R package “phyloseq” (McMurdie et al. 2013). The R package “picante” (Kembel et al. 2010) was used to compute Faith’s phylogenetic diversity of rarefied bacterial reads. Prior to compute Bray-Curtis or weighted UniFrac distances, count reads were normalized by cumulative sum scaling normalization factors (Paulson et al. 2013). To test for significantly enriched OTUs, the data were fitted to a zero-inflated Gaussian mixture model (Paulson et al. 2013). To compute Neutral and Beta-null community assembly models, R scripts were adapted for Sloan et al. 2006 and Lee et al. 2017, respectively. R scripts used in these analyses are provided via github [https://github.com/hmamine/MWDW//tree/master/community_analysis].

## Supporting information

Supplementary materiel

## Acknowledgements

We thank Janine Haueisen and Michael Habig for helpful comments to an earlier version of this manuscript. The project was funded by the DFG Collaborative Research Centerre (CRC) 1182 “Origin and Function of Metaorganisms” and the Canadian Institute for Advanced Research, CIFAR.

## Author contributions

EHS and MAH designed the experiments. MAH conducted the experiments, collected the results and analyzed the data. EZ collected *T. aestivum* seeds from the province of Kislak Antakya in Turkey. SF performed the sequencing. MAH and EHS wrote the manuscript. The authors declare no competing interests.

## List of supplementary figures and tables

Supplementary Fig. 1. The diversity of the bacterial and fungal microbiota associated to wild and domesticated wheat species.

Supplementary Fig. 2. Community α-diversity of bacteria.

Supplementary Fig. 3. Fungal communities α-diversity across wild and domesticated wheat species.

Supplementary Fig. 4. Community structure of the bacterial microbiota across wild and domesticated wheat species.

Supplementary Fig. 5. The structure of the wheat mycobiota.

Supplementary Fig. 6. Comparing the community structure of the microbiota of wild wheat to T. aestivum.

Supplementary Fig. 7. Enrichment analysis of bacteria in the phyllosphere.

Supplementary Fig. 8. Enrichment analysis of the root bacterial microbiota.

Supplementary Fig. 9. Enrichment analysis of the rhizosphere bacterial communities.

Supplementary Fig. 10. The taxonomy distribution of enrichment bacteria in wheat species.

Supplementary Fig. 11. Enrichment analysis of fungi in the phyllosphere.

Supplementary Fig. 12. Enrichment analysis of the root mycobiota.

Supplementary Fig. 13. Enrichment analysis of the rhizosphere fungal communities.

Supplementary Fig. 14. The taxonomy distribution of enrichment fungal OTUs.

Supplementary Fig. 15. Neutral model of the bacterial community assembly in wild and domesticated wheat species.

Supplementary Fig. 16. Bacterial and fungal dispersal rates.

Supplementary Fig. 17. Occurrence frequency of fungi in wild and domesticated wheat species.

Supplementary table 1. Parameters of fit of the neutral model for soil bacterial communities as seed source.

Supplementary table 2. Parameters of fit of the neutral model.

