## Supplementary materiel for "Ecological assembly processes of the bacterial and fungal microbiota of wild and domesticated wheat species"

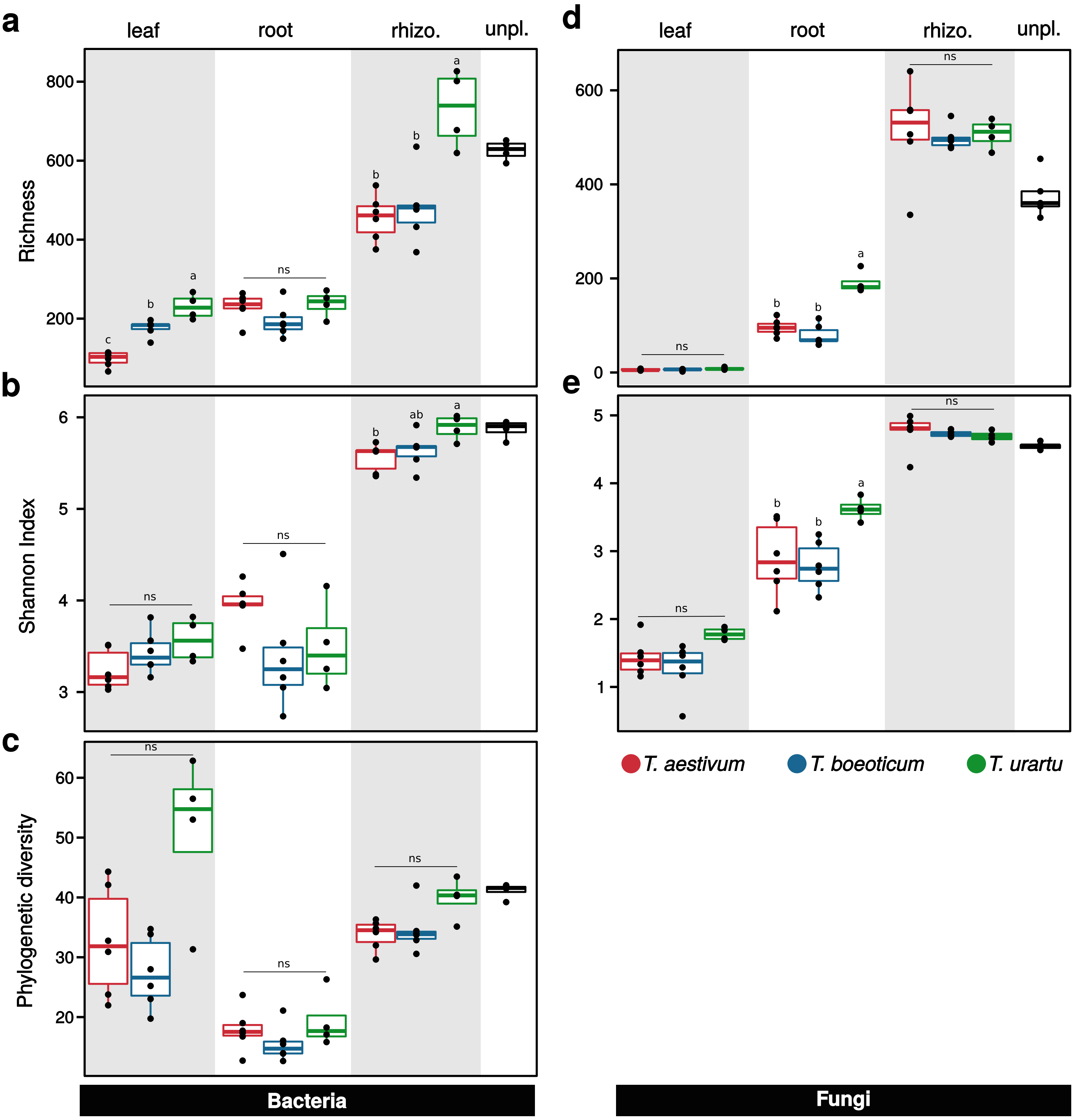


**Supplementary Fig. 1 | The diversity of the bacterial and fungal microbiota associated to wild and domesticated wheat species.** Group panels **a**, **b**, **c** and **d**, **e** show measures of community diversity of bacterial and fungal microbiota, respectively. **a**, **b** and **c**, are box-plots that show bacterial community richness, Shannon and Faith's phylogenetic indexes (rarefied to 3649 reads), respectively. Leaf and rhizosphere habitats of wild wheat *T. urartu* harbor more diverse bacterial communities than *T. boeoticum* and *T. aestivum*, these communities tend to be phylogenetically diverse. Across plant habitats, species evenness is fairly comparable between the three wheat genotypes. **d** and **e**, depict fungal community richness and Shannon index, respectively. The three wheat genotypes do not show significant differences in their mycobiota diversity, except for the roots of *T. urartu* that show increased species richness and evenness compared to *T. boeoticum* and *T. aestivum*. Color in the graph indicates the wheat genotype and black box-plot refers to unplanted soil samples. Differences between group medians were tested using Kruskal-Wallis significance test and Conover's multiple comparisons test. rhizo. And unpl. indicate rhizosphere and unplated soil samples, respectively.

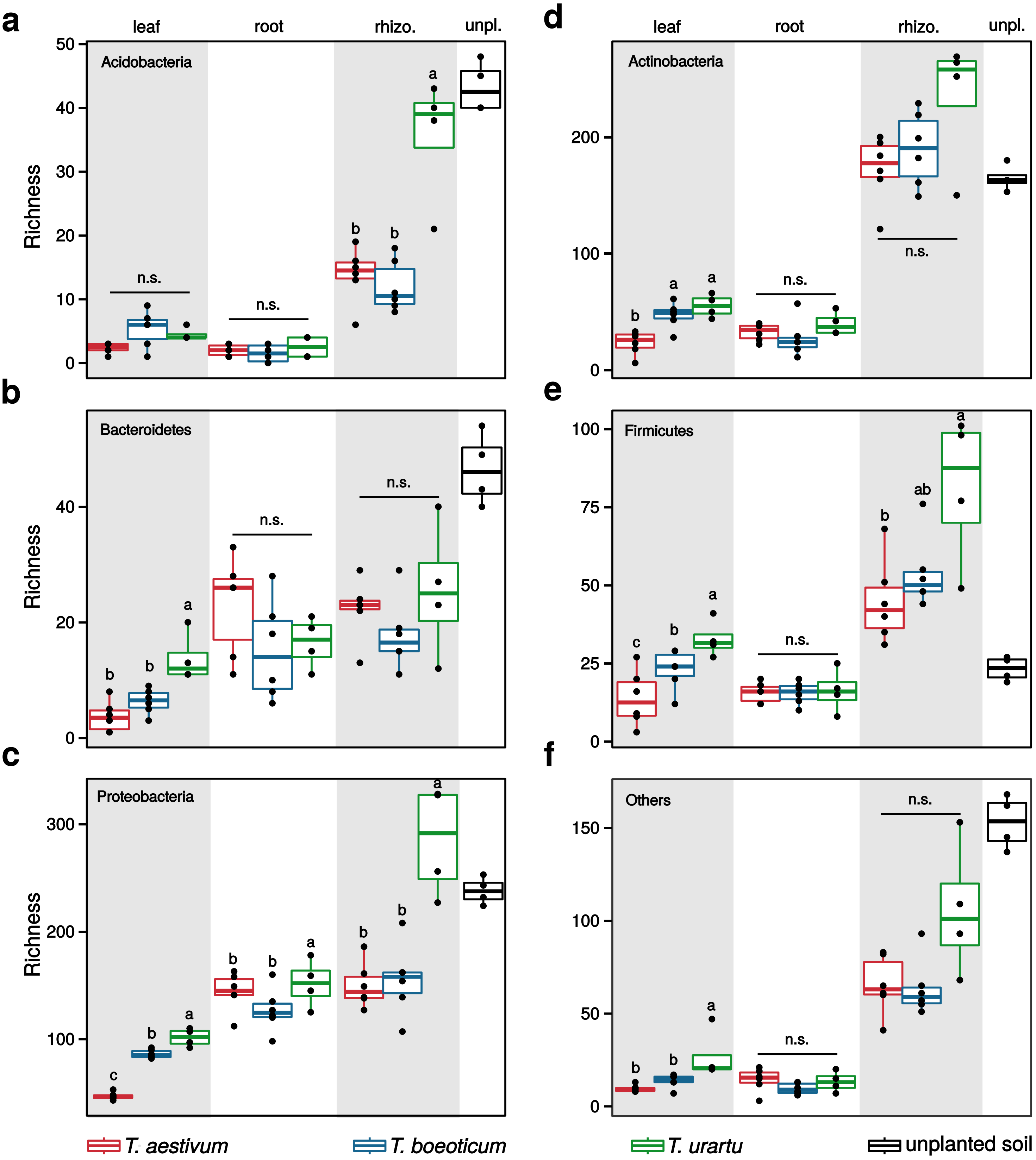


**Supplementary Fig. 2 | Community α-diversity of bacteria.** Group panels **a**, **b**, **c**, **d**, **e** and **f** show community richness across wild and domesticated wheat species for Acidobacteria, Actinoacteria, Bacteroidetes, Firmicutes, Proteobacteria and Others bacterial phyla, respectively. Reads were rarefied to an even sequencing depth of 3649 prior to subset the corresponding phylum. Colors in the graph indicate the wheat genotype and black box-plot refers to unplanted soil. Differences between group medians were tested using Kruskal-Wallis significance test Conover's multiple comparisons test. rhizo. and unpl. indicate rhizosphere and unplated soil samples, respectively.

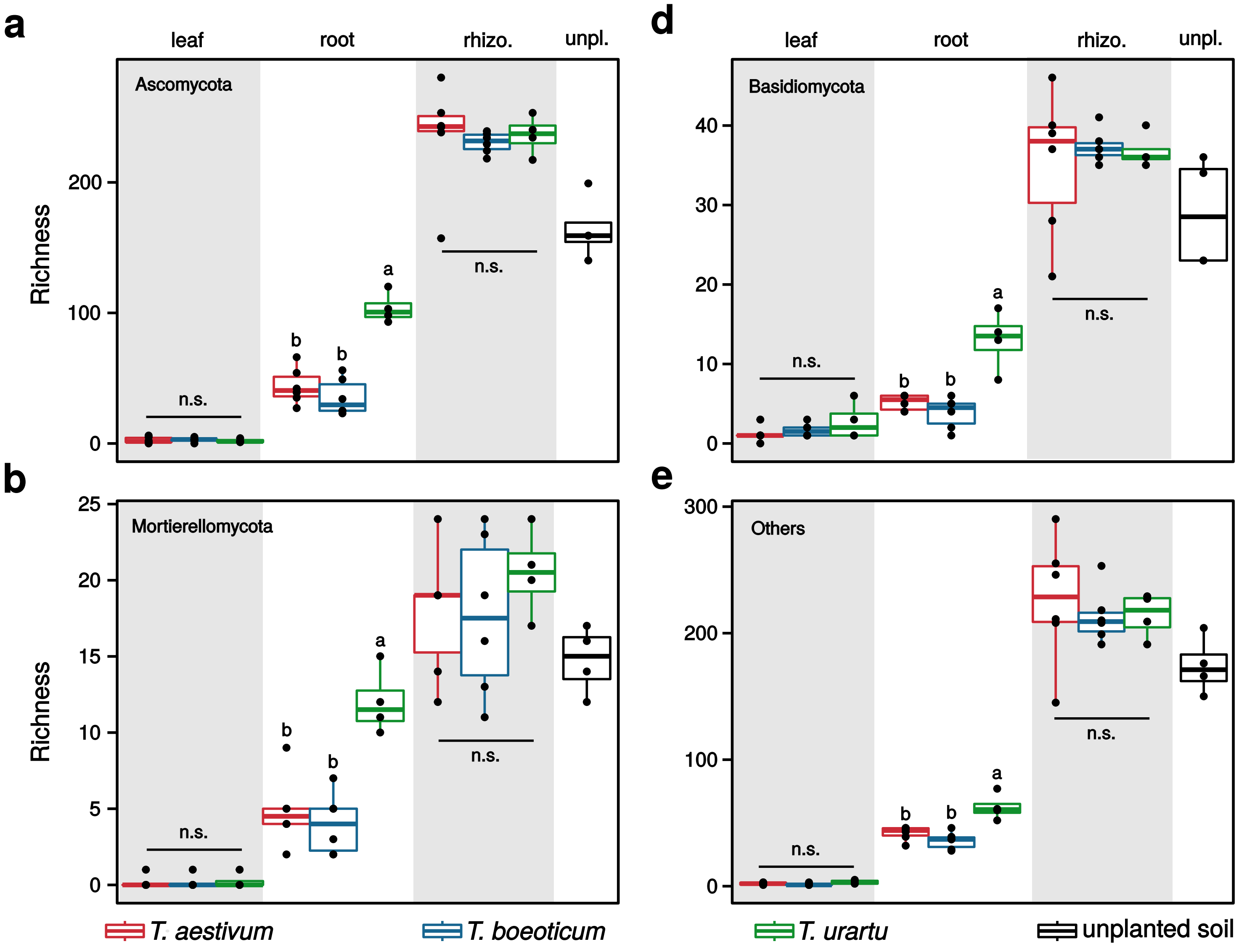


**Supplementary Fig. 3 | Fungal communities α-diversity across wild and domesticated wheat species.** Panels **a**, **b**, **c** and **d** indicate fungal species richness across wild and domesticated wheat species for Ascomcota, Basidiomycota, Mortierellomycota and others fungal phyla, respectively. Reads were not rarefied across samples prior to subset the corresponding phylum. Colors in the graph indicate the wheat genotype and black box-plot refers to unplanted soil. Differences between group medians were tested using Kruskal-Wallis significance test Conover's multiple comparisons test. rhizo. and unpl. indicate rhizosphere and unplated soil samples, respectively.

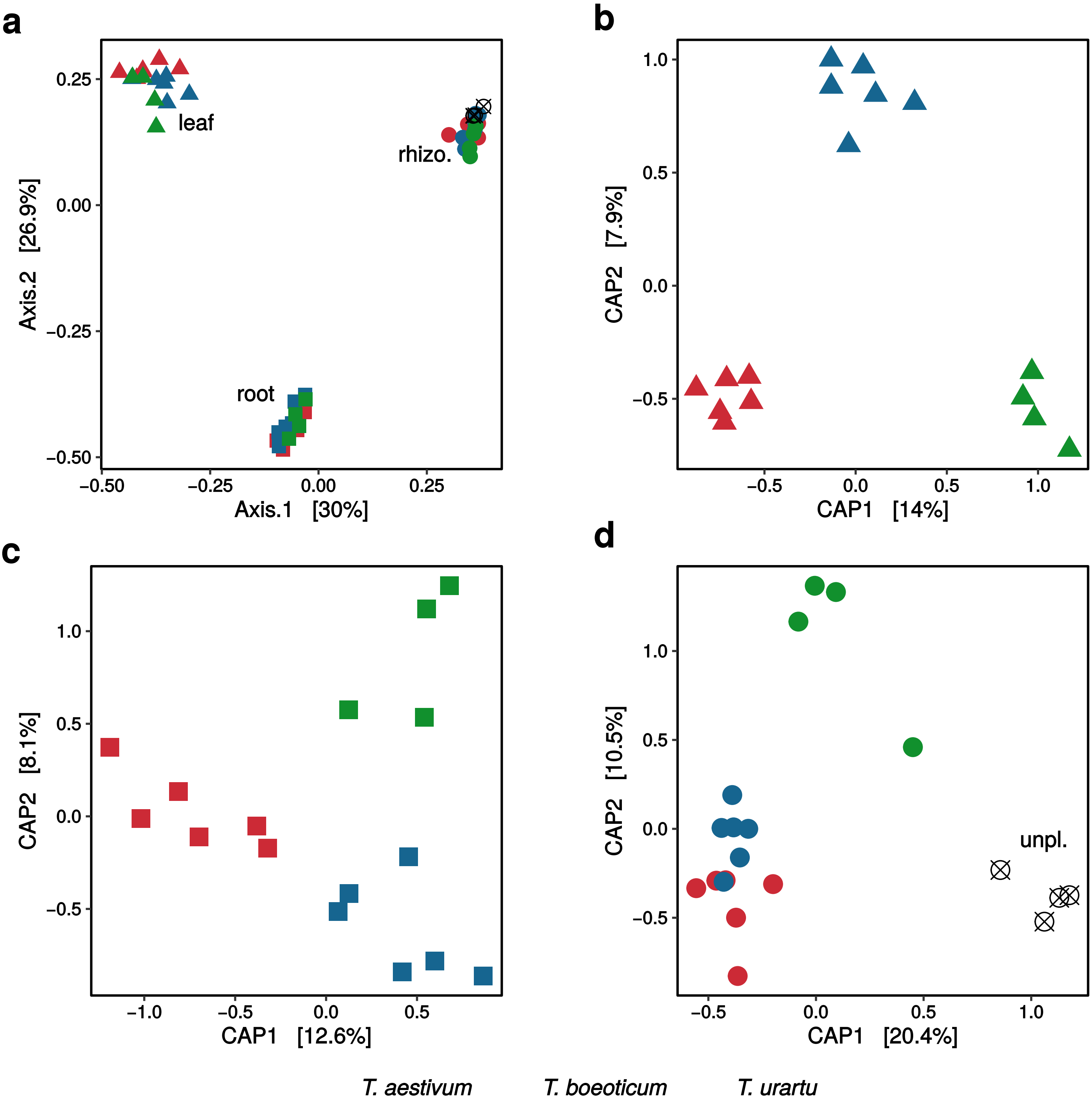


**Supplementary Fig. 4 | Community structure of the bacterial microbiota across wild and domesticated wheat species.** Panel **a**, and group panels **b**, **c** and **d** show unconstrained and constrained principal coordinates analysis (PCoA) of Bray-Curtis (BC) distances, respectively. Count reads were normalized by cumulative sum scaling normalization factors prior to compute BC distances (**Methods**). Square, triangle and circle in the graphs indicate root, leaf and rhizosphere bacterial communities, respectively. Unplanted soil is represented by a crossed unfilled and black circle. **a**, the plant habitat (i.e. leaf, root and rhizosphere) is a strong factor shaping the bacterial microbiota of wheat. The constrained analyses of BC distances indicate that the three wheat genotypes assembly distinct bacterial communities in the phyllosphere (panel **b** - $F_{2,13}$=1.82, p=0.001), roots (panel **c** - $F_{2,13}$=1.69, p=0.003) and rhizosphere (panel **d** - $F_{2,16}$=3.04, p=0.001).

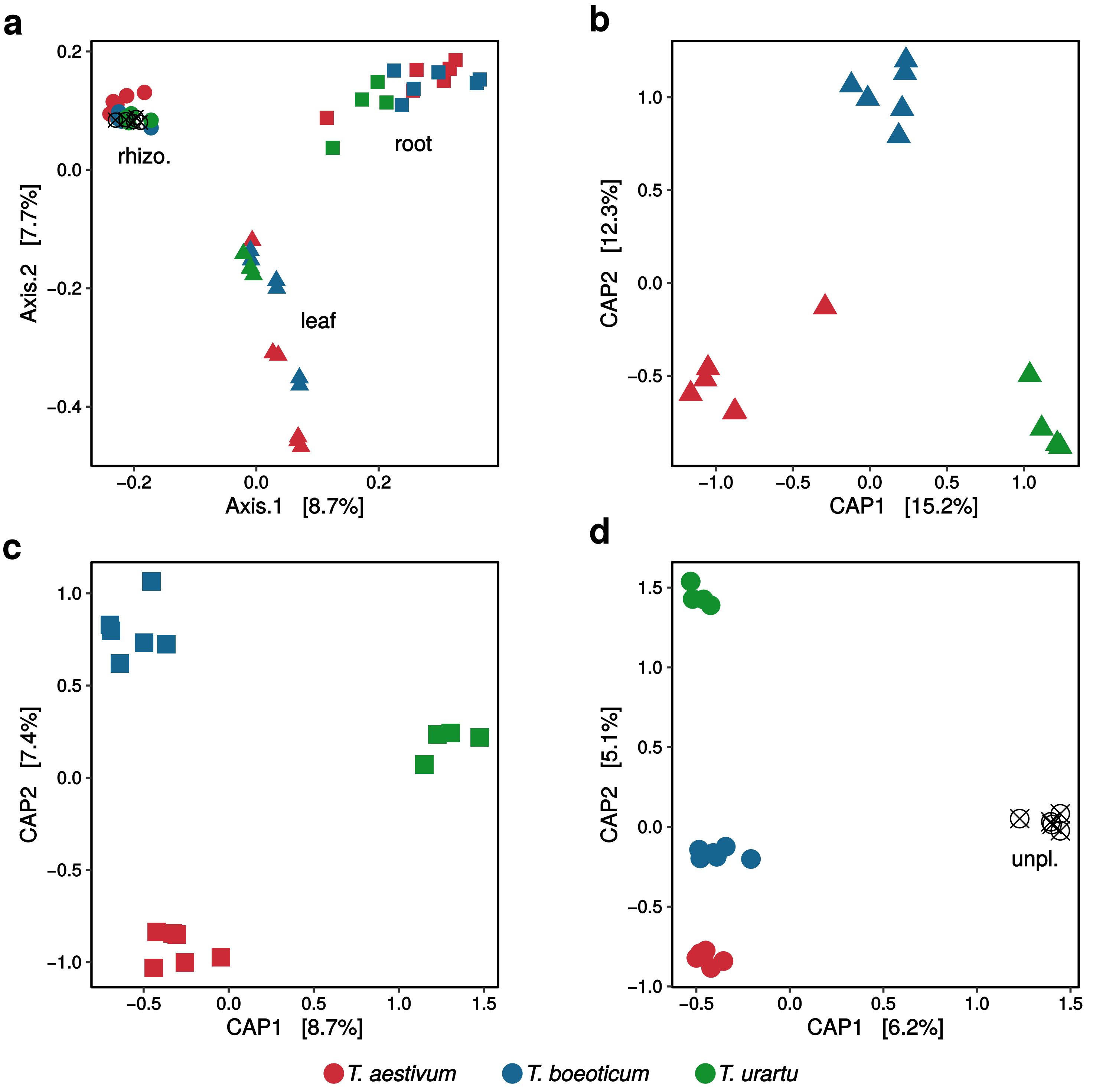


**Supplementary Fig. 5 | The structure of the wheat mycobiota.** Panel **a**, and group panels **b**, **c** and **d** show unconstrained and constrained principal coordinates analysis (PCoA) of Bray-Curtis (BC) distances, respectively. Count reads were normalized by cumulative sum scaling normalization factors prior to compute BC distances (Methods). Square, triangle and circle in the graphs indicate root, leaf and rhizosphere bacterial communities, respectively. Unplanted soil is represented by a crossed unfilled and black circle. **a**, leaf, root and rhizosphere assemble distinct mycobiota. From panels **b**, **c** and **d**, each wheat genotype assembles a distinct mycobiota in the phyllosphere ($F_{2,13}$=2.46, p=0.001), roots ($F_{2,13}$=1.24, p=0.003) and rhizosphere ($F_{2,16}$=1.08, p=0.001), respectively.

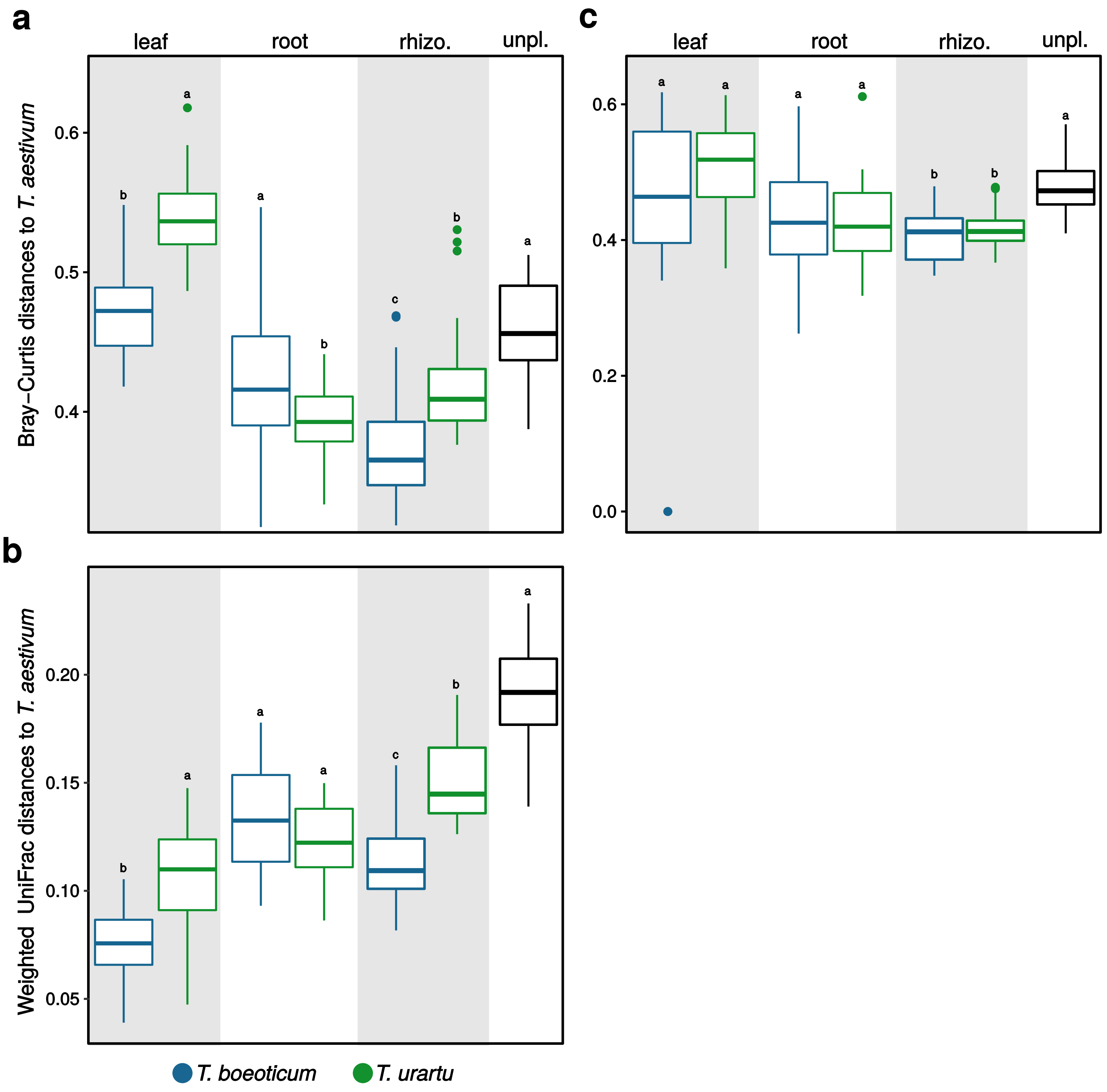


**Supplementary Fig. 6 | Comparing the community structure of the microbiota of wild wheat to *T. aestivum*.** Panel **a**, **b** depict BC and weighted UniFrac distances of bacterial communities to *T. aestivum*. The leaf and rhizosphere bacterial microbiota of *T. boeoticum* are less dissimilar in their community membership and phylogenetic compositions to *T. aestivum* than *T. urartu*. **c**, BC distances of fungal communities to *T. aestivum* do not show similar community composition phenotypes.

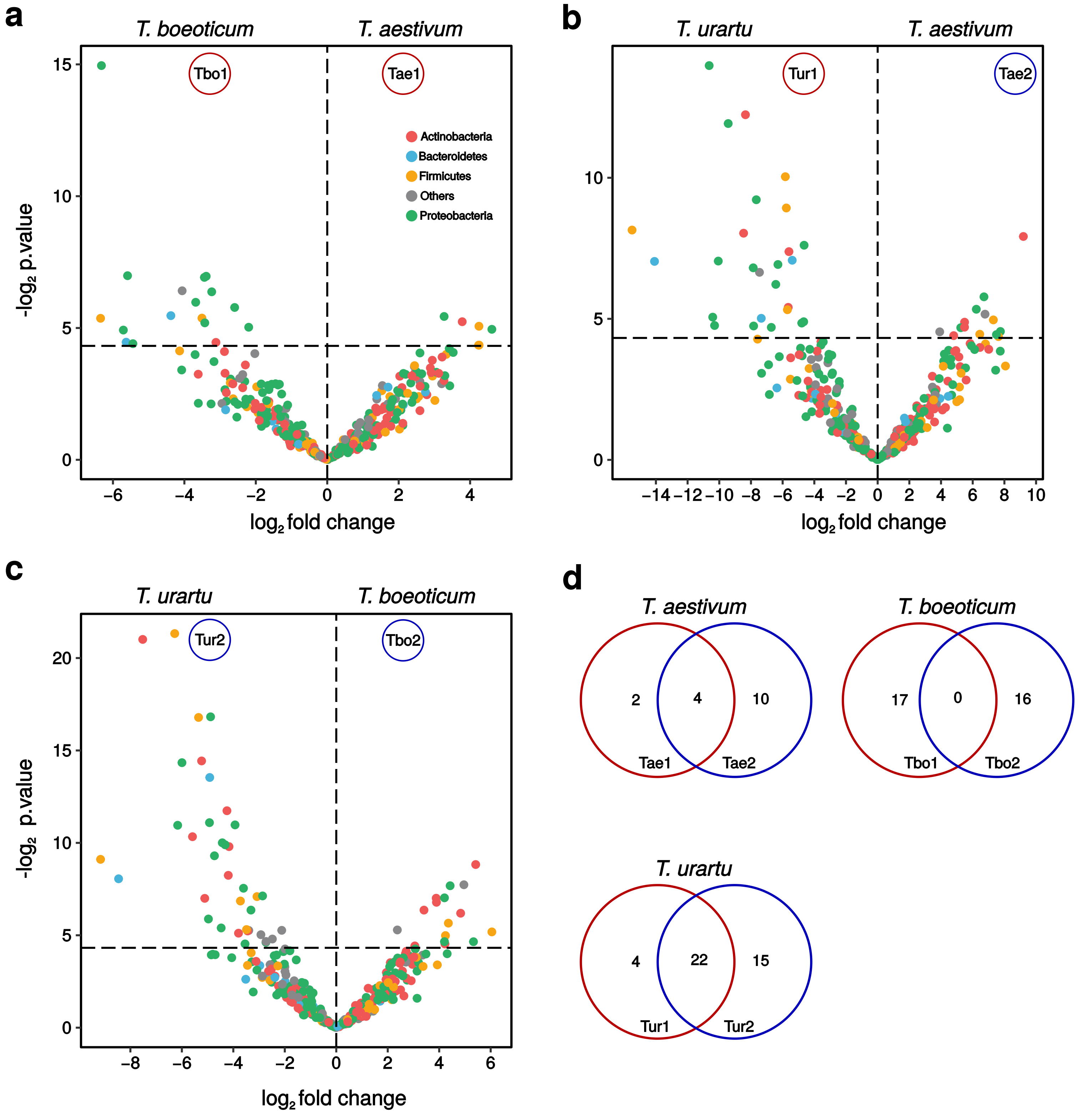


**Supplementary Fig. 7 | Enrichment analysis of bacteria in the phyllosphere**. Panel **a**, **b** and **c** show volcano plots that depict changes in the relative abundance of bacterial OTUs (x-axis) and corresponding p-value (y-axis) for the comparisons *T. aestivum* vs *T. boeoticum*, *T. aestivum* vs *T. urartu* and *T. boeoticum* vs *T. urartu*, respectively. Each circle corresponds to a bacterial OTU and the color indicates the phylum. Vertical line in the graph shows the critical p-value of 0.05. For each pairwise comparison, OTUs with an occurrence < 2 were trimmed out and count data were normalized using cumulative sum scaling factors and then fitted to zero-inflated Gaussian mixture model. Tbo1 and Tbo2 indicate OTUs that are significantly enriched in *T. boeoticum* computed from the comparison with *T. aestivum* and *T. urartu*, respectively. Tae1 and Tae2 indicate OTUs that are significantly enriched in *T. aestivum* computed from the comparison with *T. boeoticum* and *T. urartu*, respectively. Tur1 and Tur2 indicate OTUs that are significantly enriched in *T. urartu* computed from the comparison with *T. aestivum* and *T. boeoticum*, respectively. **d**, Venn diagrams depict significantly enriched (p-value < 0.05) bacterial OTUs that are unique or shared to a wheat genotype (i.e. *T. aestivum*, *T. boeoticum* or *T. urartu*) revealed by the enrichment tests in the panels a, b and c.

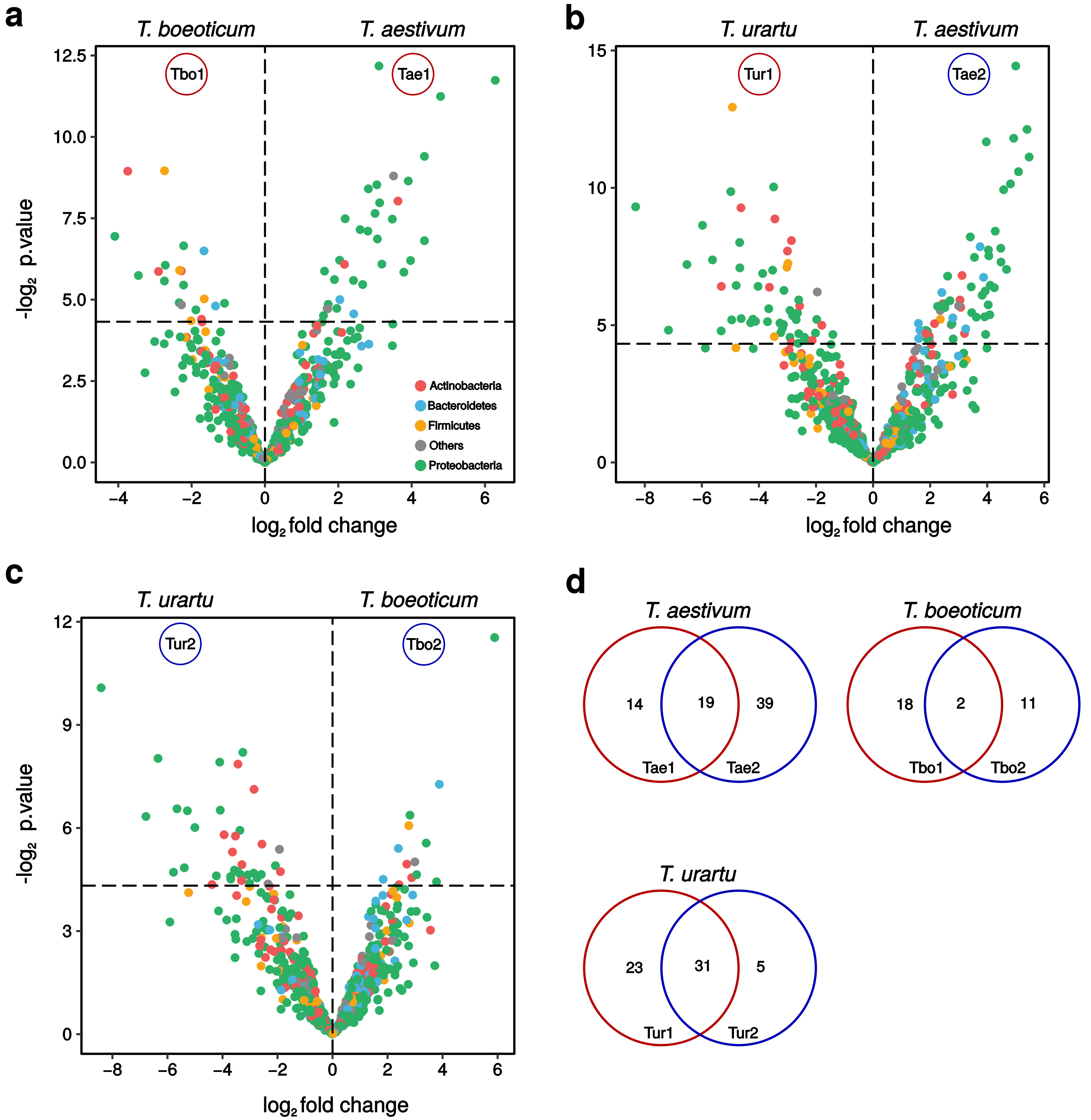


**Supplementary Fig. 8 | Enrichment analysis of the root bacterial microbiota.** Panel **a**, **b** and **c** show volcano plots for the comparisons *T. aestivum* vs *T. boeoticum*, *T. aestivum* vs *T. urartu* and *T. boeoticum* vs *T. urartu*, respectively. d, Venn diagrams depict shared and unique significantly enriched OTUs for each wheat species. Further details are indicated in the legend of supplementary figure 6.

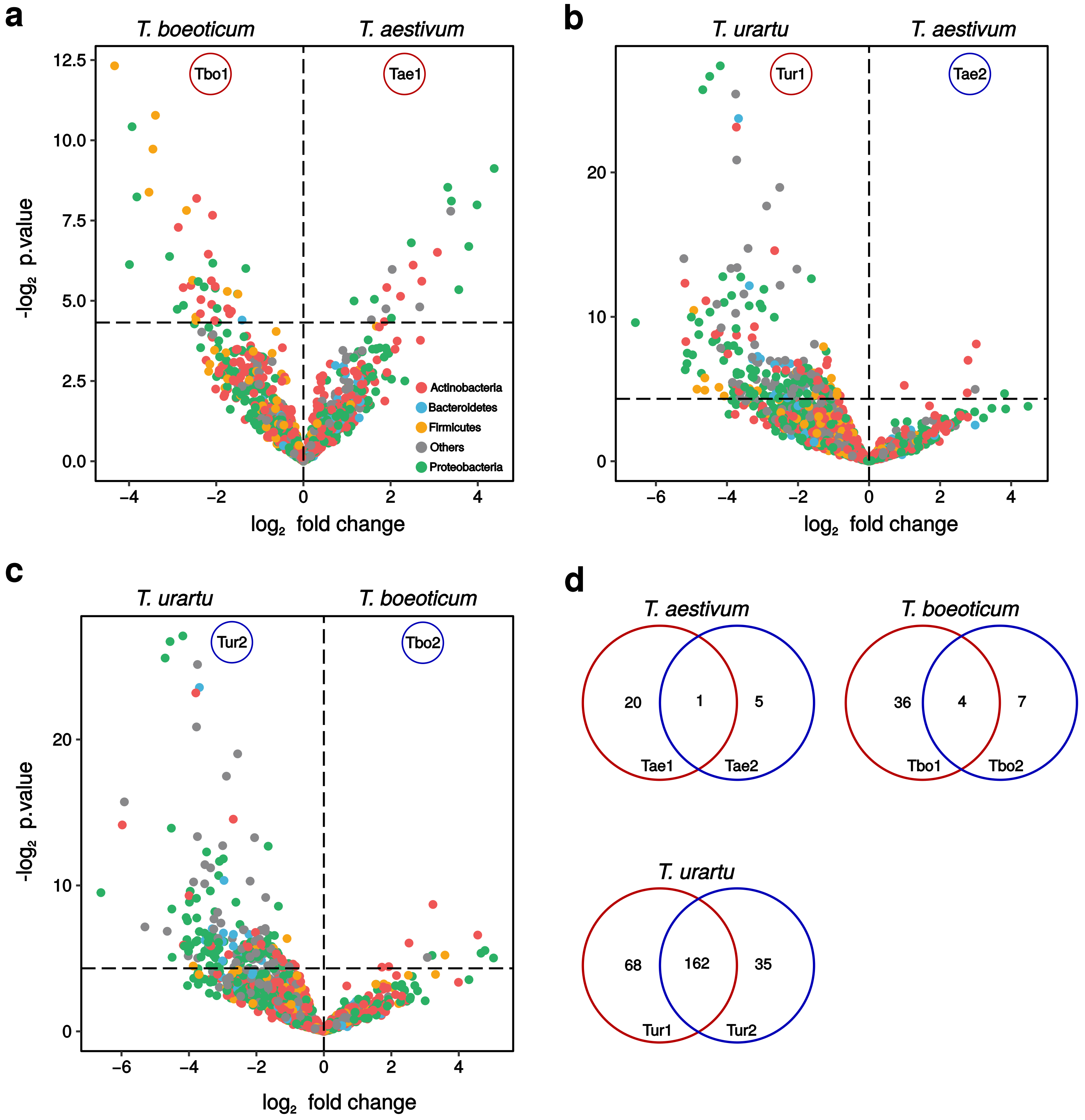


**Supplementary Fig. 9 | Enrichment analysis of the rhizosphere bacterial communities.** Panel **a**, **b** and **c** show volcano plots for the comparisons *T. aestivum* vs *T. boeoticum*, *T. aestivum* vs *T. urartu* and *T. boeoticum* vs *T. urartu*, respectively. **d**, Venn diagrams depict shared and unique significantly enriched OTUs for each wheat species. Further details are indicated in the legend of supplementary figure 6.

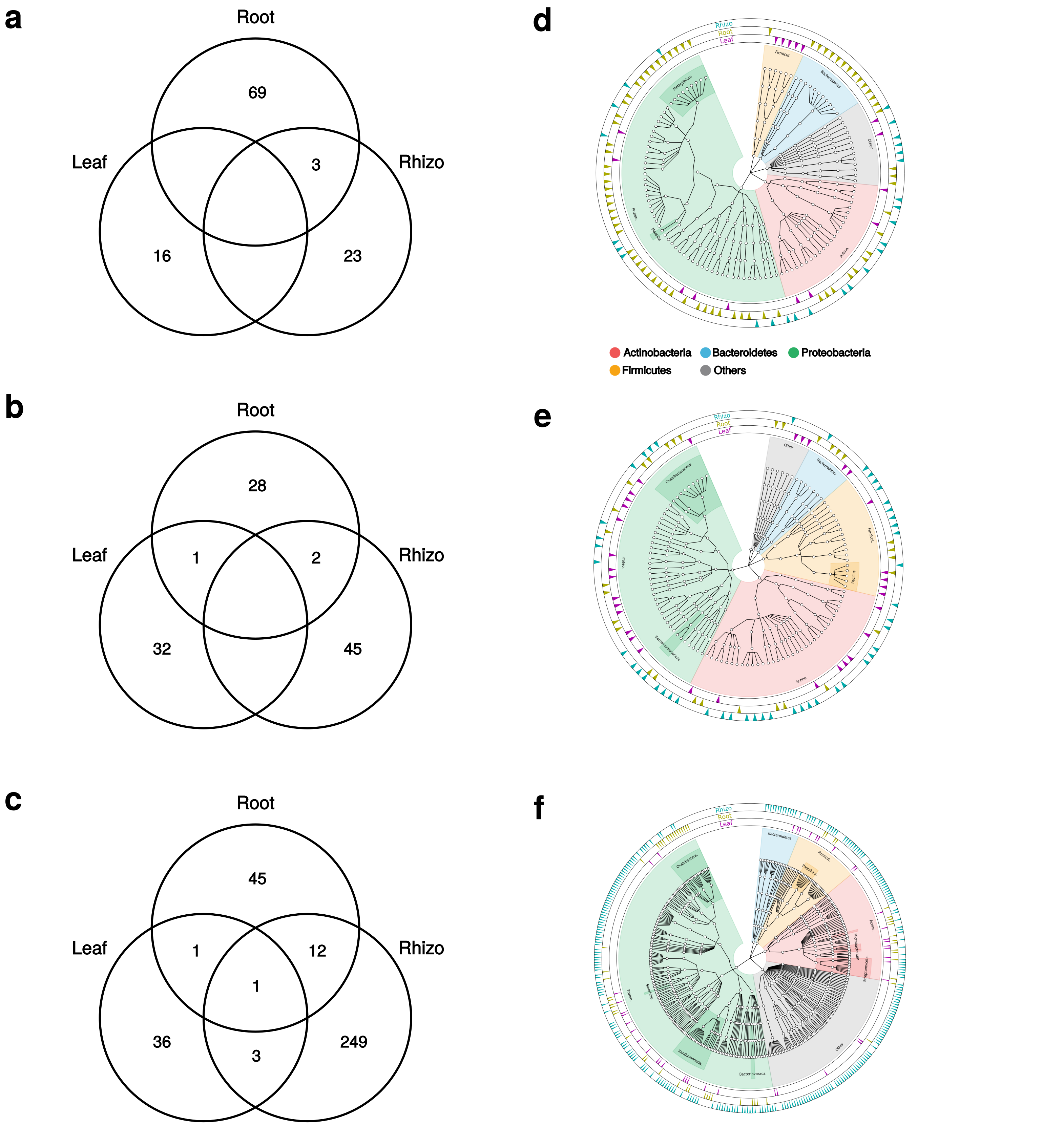


**Supplementary Fig. 10 | The taxonomy distribution of enrichment bacteria in wheat species.** Panels **a**, **b** and **c** show shared and unique bacterial OTUs enriched in the leaf (supplementary Fig 7), root (supplementary Fig 8) and rhizosphere (supplementary Fig 9) of *T. aestivum*, *T. boeoticum* and *T. urartu*, respectively. Panels **d**, **e** and **f** show taxonomic dendrograms of enriched bacterial OTUs in *T. aestivum*, *T. boeoticum* and *T. urartu*, respectively.

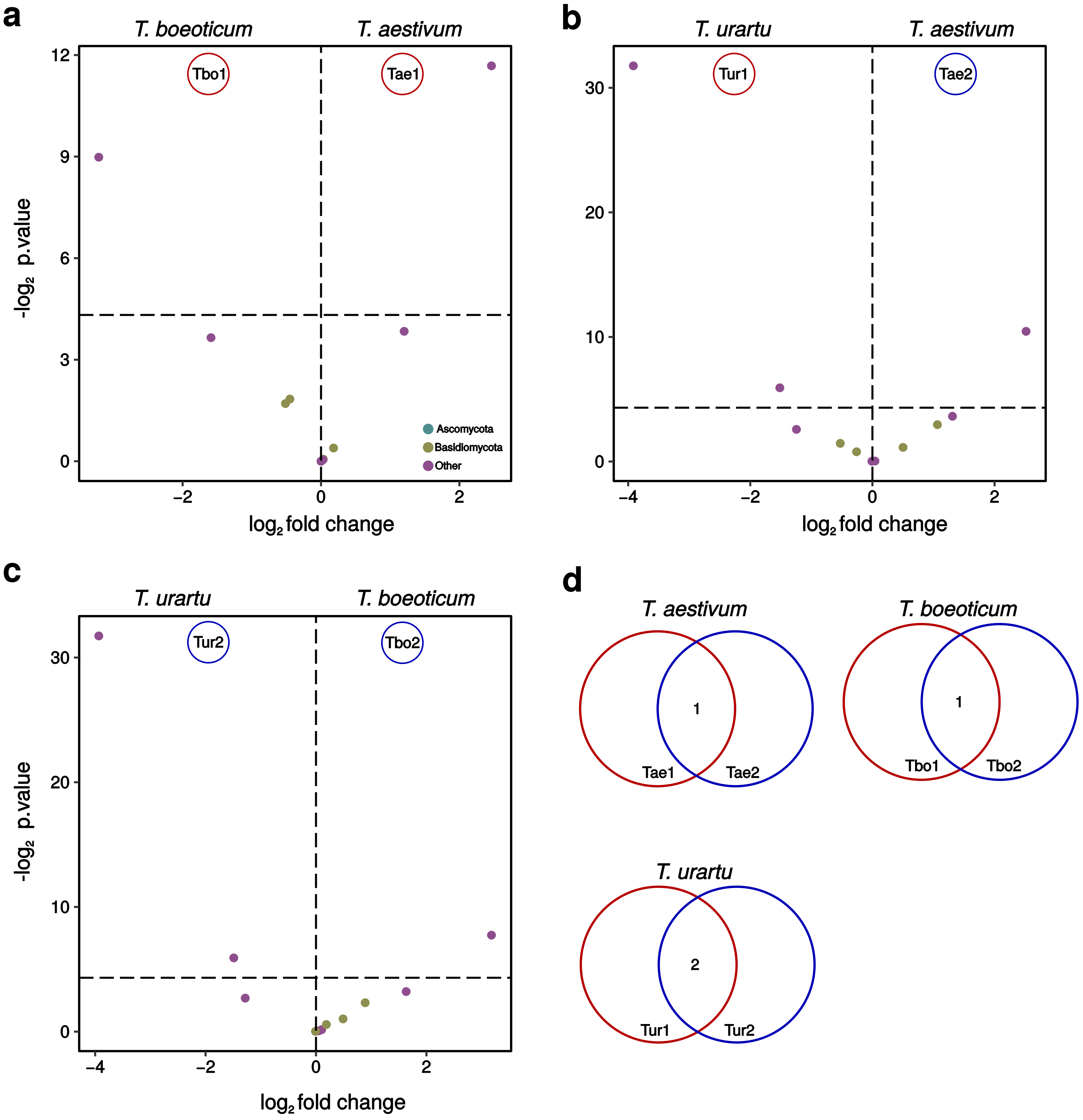


**Supplementary Fig. 11 | Enrichment analysis of fungi in the phyllosphere.** Panel **a**, **b** and **c** depict changes in the relative abundance of fungal OTUs (x-axis) and corresponding p-value (y-axis) revealed through the comparisons *T. aestivum* vs *T. boeoticum*, *T. aestivum* vs *T. urartu* and *T. boeoticum* vs *T. urartu*, respectively. Each circle corresponds to a fungal OTU and the color indicates the phylum. Vertical line in the graph shows the critical p-value of 0.05. For each pairwise comparison, OTUs with an occurrence < 2 were trimmed out and count data were normalized using cumulative sum scaling factors and then fitted to zero-inflated Gaussian mixture model. Tbo1 and Tbo2 indicate OTUs that are significantly enriched in T. boeoticum computed from the comparison with T. aestivum and T. urartu, respectively. Tae1 and Tae2 indicate OTUs that are significantly enriched in T. aestivum computed from the comparison with T. boeoticum and T. urartu, respectively. Tur1 and Tur2 indicate OTUs that are significantly enriched in T. urartu computed from the comparison with T. aestivum and T. boeoticum, respectively. **d**, Venn diagrams depict significantly enriched (p-value < 0.05) fungal OTUs that are unique or shared to a wheat genotype (i.e. *T. aestivum*, *T. boeoticum* or *T. urartu*) revealed by the enrichment tests in the panels a, b and c.

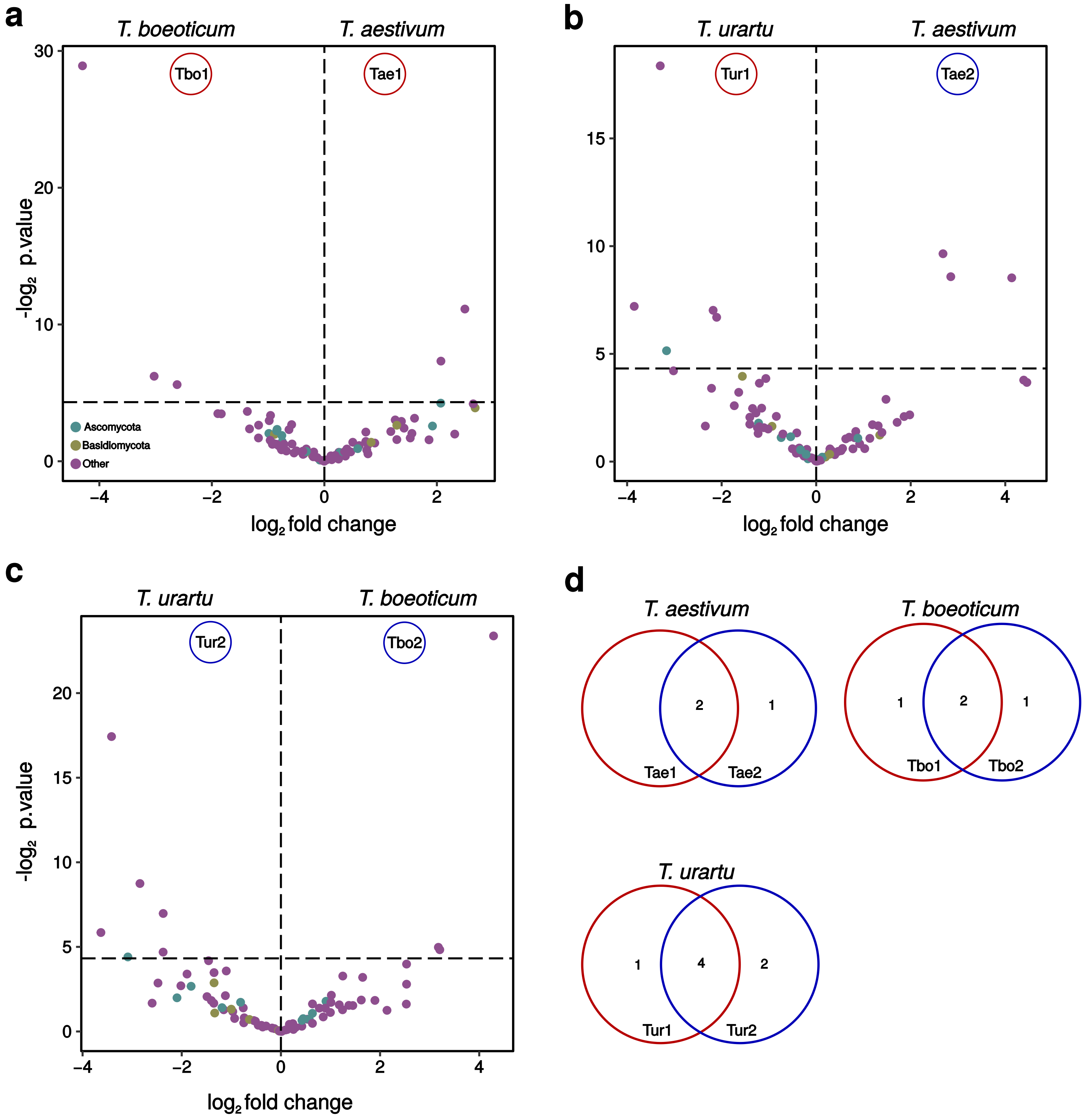


**Supplementary Fig. 12 | Enrichment analysis of the root mycobiota.** Panel **a**, **b** and **c** show volcano plots for the comparisons *T. aestivum* vs *T. boeoticum*, *T. aestivum* vs *T. urartu* and *T. boeoticum* vs *T. urartu*, respectively. d, Venn diagrams depict enriched OTUs. Further details are indicated in the legend of supplementary figure 10.

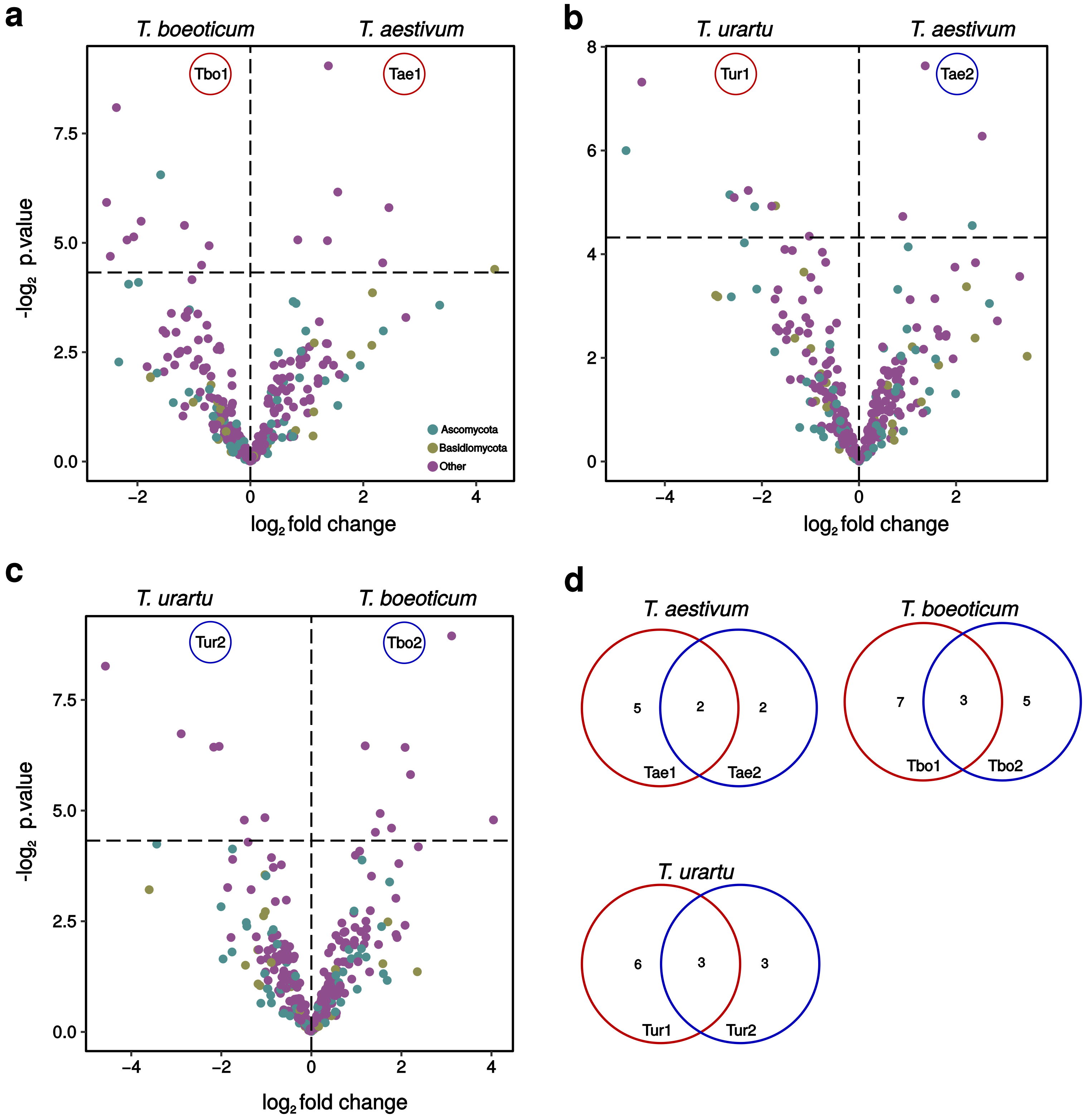


**Supplementary Fig. 13 | Enrichment analysis of the rhizosphere fungal communities.** Panel **a**, **b** and **c** show volcano plots for the comparisons *T. aestivum* vs *T. boeoticum*, *T. aestivum* vs *T. urartu* and *T. boeoticum* vs *T. urartu*, respectively. **d**, Venn diagrams depict significantly enriched fungal OTUs. Further details are indicated in the legend of supplementary figure 10.

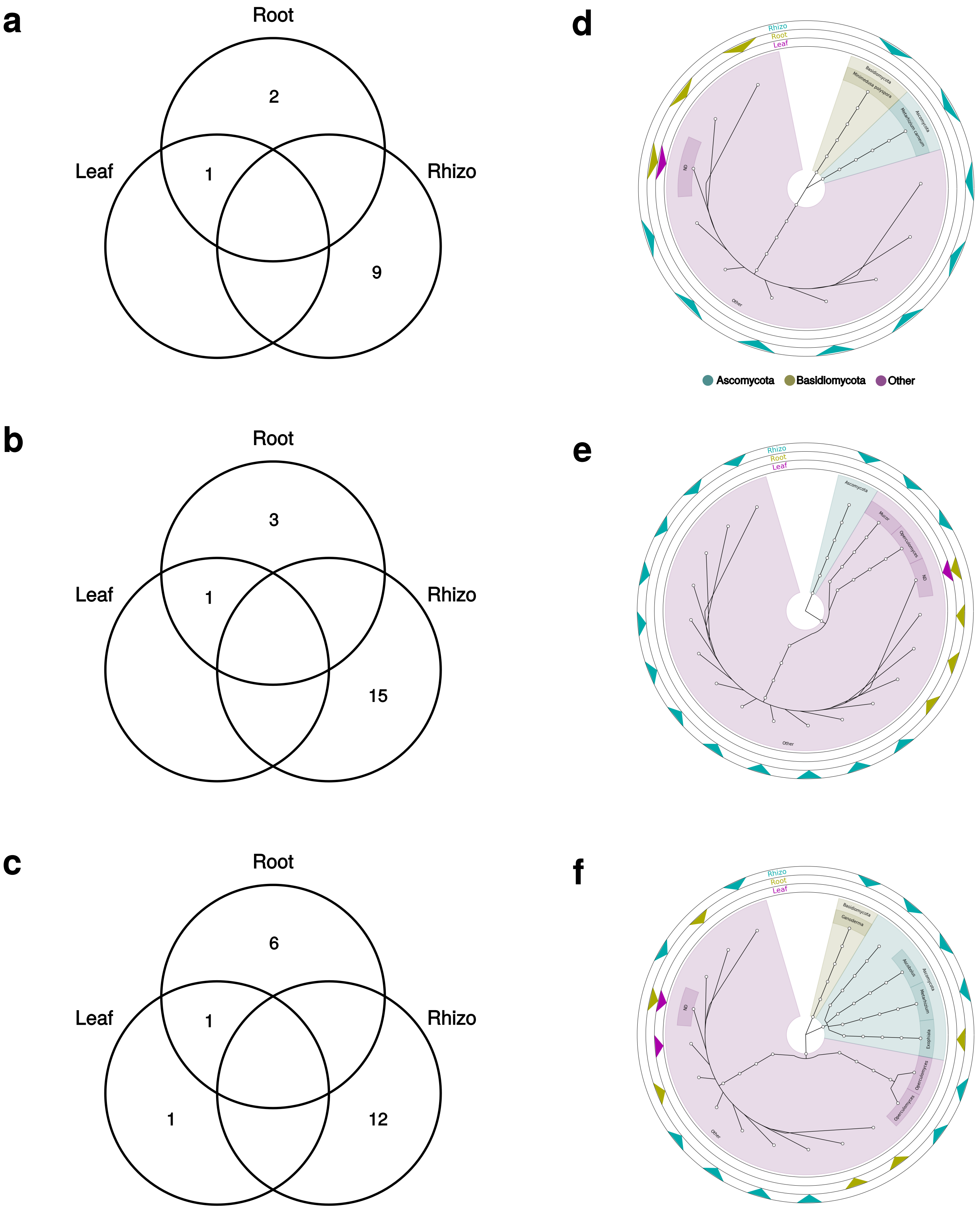


**Supplementary Fig. 14 | The taxonomy distribution of enrichment fungal OTUs.** Panels **a**, **b** and **c** show enriched OTUs in the leaf (supplementary Fig 11), root (supplementary Fig 12) and rhizosphere (supplementary Fig 13) of *T. aestivum*, *T. boeoticum* and *T. urartu*, respectively. Panels **d**, **e** and **f** show taxonomic dendrograms of enriched fungal OTUs in *T. aestivum*, *T. boeoticum* and *T. urartu*, respectively.

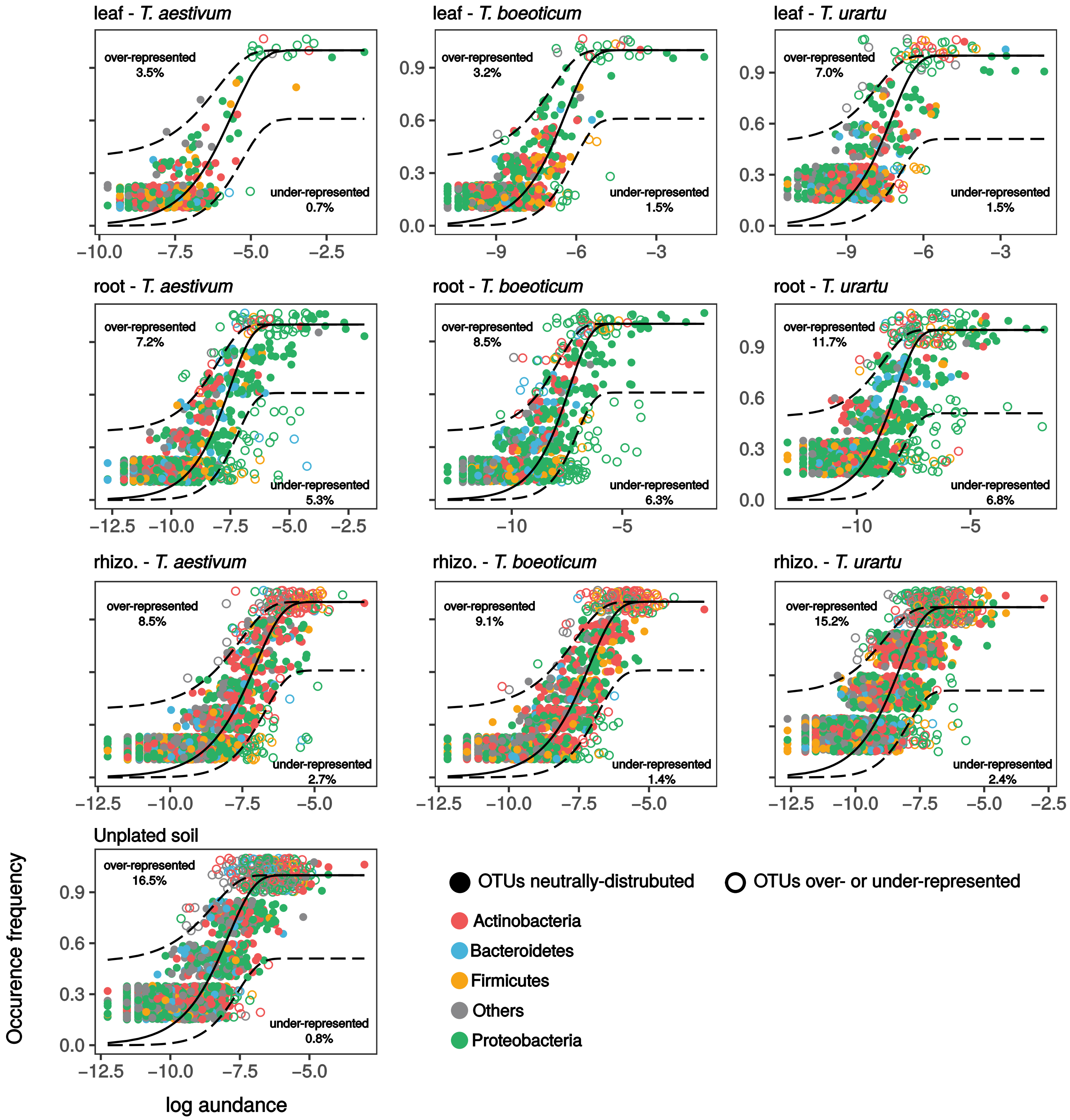


**Supplementary Fig. 15 | Neutral model of the bacterial community assembly in wild and domesticated wheat species.** Each circle indicates a bacterial OTUs. Color in the shape indicates bacterial phylum. Filled or unfilled circles indicates bacterial OTUs that follow or deviate from neutrality. Dashed lines indicate the confidence interval of 95% around the neutral prediction (continued line).

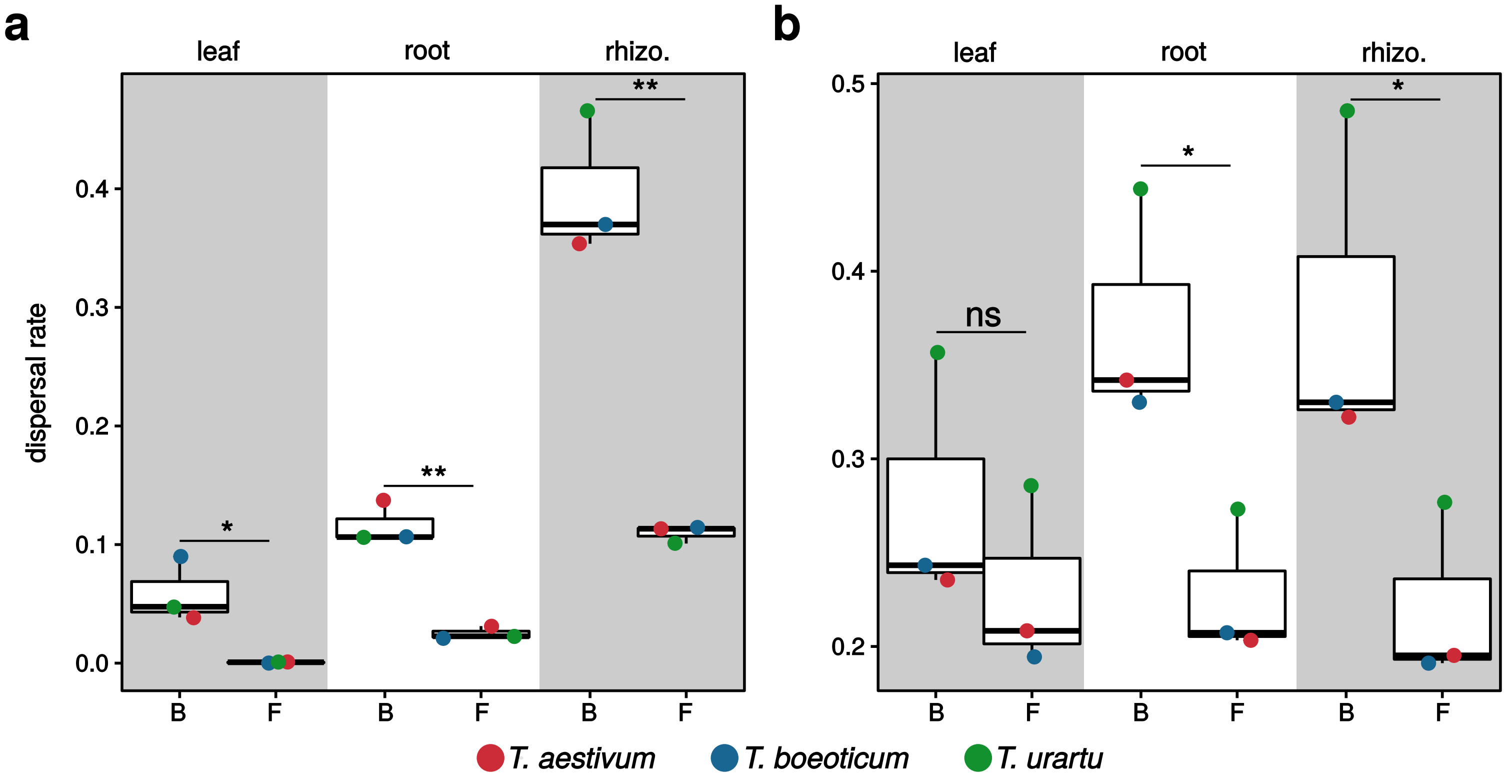


**Supplementary Fig. 16 | Bacterial and fungal dispersal rates. a**, the box-plot depict dispersal rates of bacteria and fungi from the soil. The dispersal rates from soil corresponds to percentage of shared OTUs between the habitat leaf, root or rhizosphere and the unplanted soil. **b**, the box-plots depict within-group dispersal rates of bacteria and fungi. The within-group dispersal rate corresponds to the sum of OTUs occurrence frequencies across replicates divided by the sum of observed OTUs within these replicates. B and F indicate in the graphs bacteria and fungi, respectively. Color in the circle indicates wheat genotype. Test for significance using t-Test. ns; p-value > 0.05, *; p-value <0.05, **; p-value < 0.01.

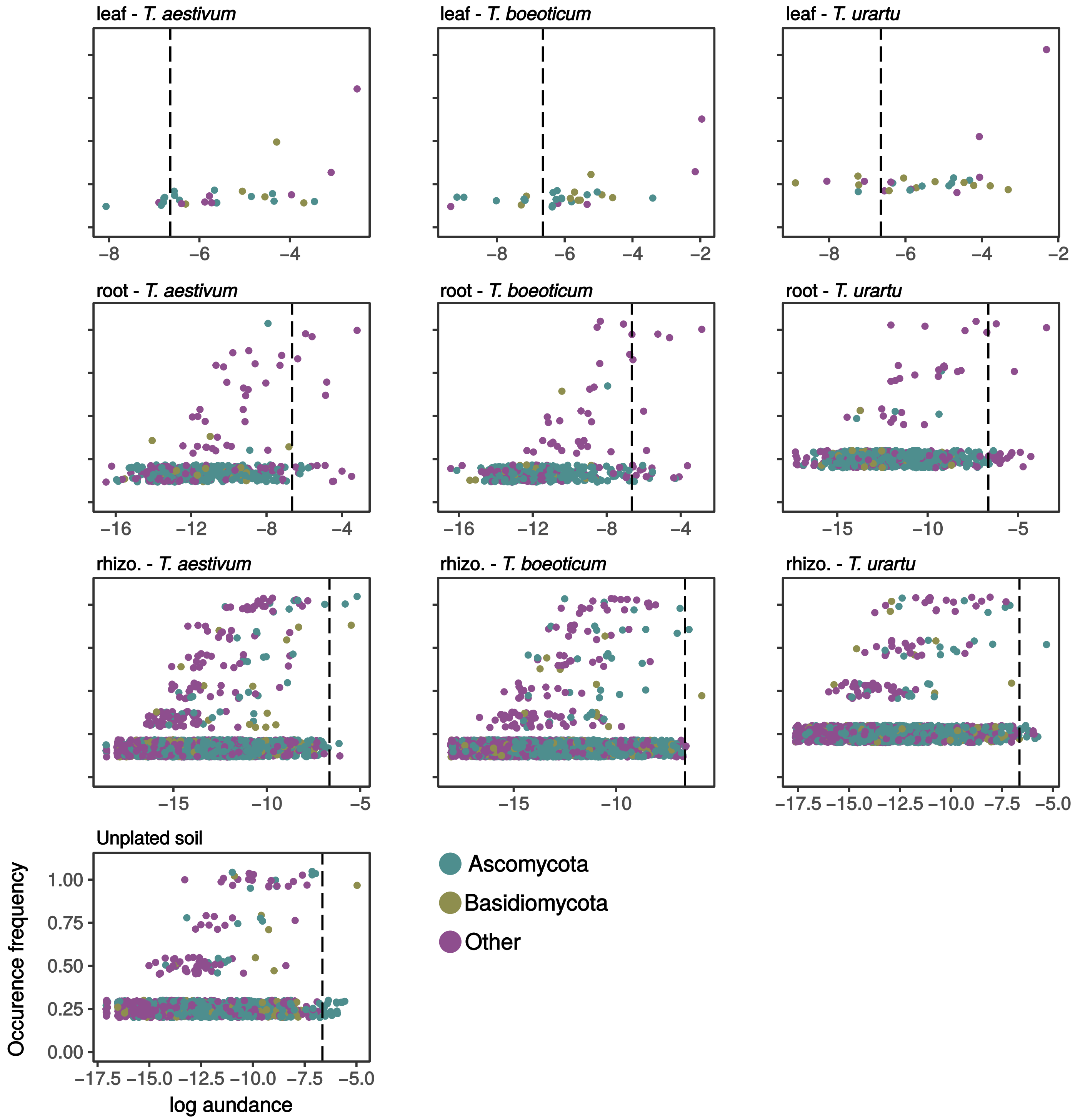


**Supplementary Fig. 17 | Occurrence frequency of fungi in wild and domesticated wheat species.** The plots show the occurrence frequency (y-axis) and log relative abundance (x-axis) of fungal OTUs in the leaf, root and rhizosphere habitats of wild and domesticated wheat species. Each circle corresponds to a fungal OTUs and color indicates the phylum. Vertical dashed line indicates the 1 % relative abundance.

**Supplementary table 1 | Parameters of fit of the neutral model for soil bacterial communities as seed source.** The goodness-of-fit of the Sloan model is indicated by (Rsqr) and computed immigration coefficient to the model is indicated by (m). (RSME) indicates the root-mean-square error.

| | | m | Rsqr | RMSE | N | Samples | Richness |
|---|---|---|---|---|---|---|---|
| leaf | Tae | 0.00010 | -0.37524 | 0.17631 | 5583.50 | 6 | 69 |
| leaf | Tbo | 0.00019 | -0.77577 | 0.18341 | 16649.33 | 6 | 161 |
| leaf | Tur | 0.00034 | -0.90012 | 0.33047 | 39868.25 | 4 | 85 |
| root | Tae | 0.00055 | -0.39393 | 0.33822 | 54346.00 | 6 | 245 |
| root | Tbo | 0.00067 | -0.51387 | 0.37654 | 67019.17 | 6 | 190 |
| root | Tur | 0.00223 | -0.85750 | 0.43460 | 116710.75 | 4 | 190 |
| rhizo. | Tae | 0.00353 | 0.12659 | 0.29657 | 31977.00 | 6 | 632 |
| rhizo. | Tbo | 0.00366 | 0.07768 | 0.30643 | 32660.50 | 6 | 661 |
| rhizo. | Tur | 0.01873 | -0.07413 | 0.30535 | 77046.00 | 4 | 832 |

**Supplementary table 2 | Parameters of fit of the Neutral model.** The goodness-of-fit of the Sloan model is indicated by (Rsqr) and computed immigration coefficient to the model is indicated by (m). (AIC.sncm) and (BIC.sncm) indicate Akaike and Bayesian information criterion of the fit, respectively. (RSME) and (m.ci) indicate the root-mean-square error and confidence interval of the model fit, respectively.

| | | m | m.ci | sncm.LL | Rsqr | RMSE | AIC.sncm | BIC.sncm | N | Samples | Richness | Detect |
|---|---|---|---|---|---|---|---|---|---|---|---|---|
| **leaf** | Tae | 0.01236 | 0.00122 | -289.49508 | 0.57818 | 0.12239 | -574.99016 | -566.89070 | 5583.50 | 6 | 424 | 0.00018 |
| | Tbo | 0.00839 | 0.00065 | -454.49447 | 0.46340 | 0.13530 | -904.98893 | -895.66778 | 16649.33 | 6 | 781 | 0.00006 |
| | Tur | 0.00789 | 0.00067 | -229.49720 | 0.35413 | 0.17979 | -454.99440 | -445.69902 | 39868.25 | 4 | 771 | 0.00003 |
| **root** | Tae | 0.00775 | 0.00085 | -177.68682 | 0.46968 | 0.19815 | -351.37365 | -341.79796 | 54346.00 | 6 | 887 | 0.00002 |
| | Tbo | 0.00496 | 0.00058 | -95.13520 | 0.30891 | 0.21616 | -186.27040 | -176.80598 | 67019.17 | 6 | 839 | 0.00001 |
| | Tur | 0.00760 | 0.00099 | 27.66767 | 0.17678 | 0.25029 | 59.33535 | 68.78541 | 116710.75 | 4 | 833 | 0.00001 |
| **rhizo.** | Tae | 0.00835 | 0.00046 | -718.05272 | 0.65257 | 0.15655 | -1432.10544 | -1421.29080 | 31977.00 | 6 | 1648 | 0.00003 |
| | Tbo | 0.00889 | 0.00044 | -852.52726 | 0.70205 | 0.14727 | -1701.05453 | -1690.15902 | 32660.50 | 6 | 1716 | 0.00003 |
| | Tur | 0.01330 | 0.00076 | -430.36723 | 0.52369 | 0.19724 | -856.73446 | -845.43222 | 77046.00 | 4 | 2103 | 0.00001 |
| | Unpl. | 0.01355 | 0.00075 | -551.36359 | 0.62543 | 0.17778 | -1098.72718 | -1087.75059 | 52466.50 | 4 | 1787 | 0.00002 |
